# When and why an essential popular mineral—calcium—is an effective antifungal adjuvant against the human pathogen?

**DOI:** 10.1101/2020.08.23.263939

**Authors:** Chi Zhang, Yiran Ren, Huiyu Gu, Lu Gao, Yuanwei Zhang, Ling Lu

## Abstract

In eukaryotes, calcium not only is an essential mineral nutrient but also serves as an intracellular second messenger that is necessary for many physiological processes. Here, we show that exogenous calcium is toxic when fungal cells lack functional calcineurin, a calcium-dependent protein phosphatase that acts as the central regulator of the calcium signaling pathway. By monitoring intracellular calcium, particularly by tracking vacuolar calcium dynamics in living cells through a novel procedure using modified aequorin, we found that calcineurin dysfunction perturbed calcium homeostasis in intracellular compartments including the cytosol, mitochondria, and vacuole, leading to drastic autophagy global organelle fragmentation, and even lastly resulting in cell death upon an extracellular calcium stimulus. Notably, the defective phenotypes seen with calcineurin mutants can be significantly suppressed by alleviating a cytosolic calcium overload or increasing vacuolar calcium storage capacity, suggesting toxicity of exogenous calcium to calcineurin mutants is tightly associated with abnormal cytosolic calcium accumulation and vacuolar calcium storage capacity deficiency. Our findings provide insights into how calcineurin regulates intracellular calcium homeostasis for cell survival and may have important implications for antifungal therapy and clinical drug administration.

## Introduction

Calcium (Ca^2+^), as an intracellular second messenger in eukaryotic cells, plays a critical role in the regulation of various cellular processes including synaptic transmission, secretion, and cytokinesis (*Islam, 2020*; *Rosendo-Pineda, Moreno, & Vaca, 2020*). The intracellular Ca^2+^ concentration is strictly and precisely controlled by a sophisticated calcium homeostasis system, which is composed of various calcium channels, calcium pumps, and calcium antiporters (*Carafoli, 1987*; *Cyert & Philpott, 2013*; *Rosendo-Pineda et al., 2020*). Overwhelming evidence demonstrates that calcineurin, the conserved Ca^2+^-calmodulin (CaM) activated protein phosphatase, is a master regulator of intracellular calcium homeostasis in fungi (*Blatzer & Latge, 2017*). Calcineurin is required for the regulation of cation homeostasis, morphogenesis, cell-wall integrity, and pathogenesis (*H. S. Park, Lee, Cardenas, & Heitman, 2019*). However, the mechanism by which calcineurin regulates the optimal Ca^2+^ concentrations in the cytosol and in intracellular compartments such as the vacuole and mitochondria remain elusive.

When fungal cells encounter environmental stresses, a transient elevation of cytosolic calcium is induced primarily through calcium influx systems in the plasma membrane such as the high-affinity calcium uptake Cch1-Mid1 channel complex system (*Cyert & Philpott, 2013*; *Thewes, 2014*). Calcium-activated calcineurin then dephosphorylates the transcription factor Crz1, which induces its translocation to the nucleus to regulate gene expression. It has been shown that the free cytosolic calcium concentration ([Ca^2+^]_c_) must be maintained at a low, steady level within a narrow physiological range (50-200 nM) after a sudden increase, peaking within seconds and then sharply decreasing within 30 seconds as the excess cytosolic calcium is transiently transferred to calcium stores including the endoplasmic reticulum (ER), vacuoles, and mitochondria by various calcium pumps and calcium antiporters (*Cunningham, 2011*; *Cyert & Philpott, 2013*; *Miseta, Fu, Kellermayer, Buckley, & Bedwell, 1999*; *Miseta, Kellermayer, Aiello, Fu, & Bedwell, 1999*; *Munoz et al., 2014*). In mammals, the ER, mitochondria, and lysosomes are physically and/or functionally linked (*Dolgin, 2019*). The role of Ca^2+^ signaling in mitochondria is to determine cellular survival by controlling basal mitochondrial bioenergetics and by regulating apoptosis. Moreover, the lysosomes can act as a Ca^2+^ store for Ca^2+^ release into the cytosol, thereby influencing Ca^2+^ homeostasis. Ca^2+^ signaling in lysosomes may regulate autophagy to respond to cell damage and cell stress (*La Rovere, Roest, Bultynck, & Parys, 2016*). As a topological equivalent, the vacuole is the primary Ca^2+^ storage organelle in fungi, containing more than 90% of total cellular Ca^2+^ (*Dunn, Gable, & Beeler, 1994*). Notably, until now there was no reliable system to probe or detect the calcium concentration in fungal vacuole, probably due to its low internal pH. Vacuolar Ca^2+^ sequestration is performed by the collaboration of Ca^2+^/H^+^ exchangers in the Vcx family and the P-type ATPase Pmc family (*Forster & Kane, 2000*; *Kmetzsch et al., 2010a*). In contrast, vacuolar Ca^2+^ can be released into the cytosol through the mechanosensitive channel Yvc1 and/or transient receptor protein (TRP) channels (*Chang, Schlenstedt, Flockerzi, & Beck, 2010*; *Denis & Cyert, 2002*; *Hamamoto, Mori, Yabe, & Uozumi, 2018*; *Lange et al., 2009*; *Thewes, 2014*; *Yu et al., 2014*). Several lines of evidence in yeast-like and filamentous fungi indicate that the transcription of Pmc family members is dependent on an activated calcineurin-Crz1/CrzA complex, whereas Vcx1 activity is negatively regulated by the calcineurin complex (*Cunningham & Fink, 1996*; *Dinamarco et al., 2012*; *Ferreira et al., 2012*; *Kmetzsch et al., 2010b*; *Thewes, 2014*).

In addition to being a calcium store, the fungal vacuole is responsible for the last step of autophagy. Autophagy is a highly conserved catabolic process that mediates the self-degradation of intracellular material, such as proteins, lipids, or even entire organelles. In the conserved autophagy pathway, the double-membrane autophagosome (AP) engulfs cellular components to be delivered for degradation in the lysosome/vacuole (*Klionsky & Emr, 2000*; *Nakatogawa, 2020*). Autophagy exists at a basal level in all living cells, but it can be induced in response to various factors such as nitrogen starvation, oxidative stress, ER stress, and rapamycin drug treatment (*Liu et al., 2014*; Y. *Wang & Zhang, 2019*; *Yi, Tong, & Yu, 2018*). In most of these situations, autophagy has both beneficial and harmful effects. It can function as a protective mechanism to promote cell survival in nutritional starvation and other stresses, but excessive autophagy also leads to damage to organelles and even cell death (*Scarlatti, Granata, Meijer, & Codogno, 2009*; *White & DiPaola, 2009*) .

*Aspergillus fumigatus* is an opportunistic filamentous fungus that causes a wide spectrum of life-threatening diseases with poor treatment outcomes in immunocompromised individuals (*Denning & Bromley, 2015*). Calcineurin is considered a potential drug target due to its essential roles in fungal growth and virulence (*Juvvadi, Lee, Heitman, & Steinbach, 2017*; *Steinbach et al., 2006*; *Steinbach et al., 2007*). In this study, we observed the phenomenon that a relatively low concentration of calcium is toxic to calcineurin-mutant fungi. By monitoring the calcium dynamics in the cytosol and intracellular calcium stores with modified forms of aequorin as a calcium reporter, we found that the calcineurin mutant exhibited abnormal calcium homeostasis and triggered autophagy and global fragmentation of the nucleus and organelles under a calcium stimulus. Additionally, the defective phenotypes of the calcineurin mutant can be significantly suppressed by alleviating cytosolic calcium overload through deletion of the Ca^2+^ channel *cchA* or by increasing vacuolar calcium storage capacity with overexpressed vacuolar P-type Ca^2+^-ATPase PmcA, highlighting the feedback relationship between calcineurin and CchA and the critical role of vacuoles in the detoxification of excess calcium for fungal survival. Our findings shed light on the underlying mechanism by which calcineurin regulates calcium homeostasis for cell survival, and these results provide a rationale for combination therapy of calcium and calcineurin inhibitors to combat human fungal pathogens. Additionally, the potential toxicity to mammalian cells of combined calcium supplements and calcineurin inhibitors should be taken into consideration when administering immunosuppressive drugs to patients in clinical settings.

## Results

### Extracellular calcium is toxic to *A. fumigatus* calcineurin mutants

Calcineurin is a highly conserved Ca^2+^-calmodulin-dependent protein phosphatase that plays a central role in morphological development and cation homeostasis in fungi (*Fox & Heitman, 2002*; *H. S. Park et al., 2016*). To further characterize the functions of calcineurin in Ca^2+^ signaling transduction, we constructed deletion mutants of the catalytic subunit CnaA and the regulatory subunit CnaB by CRISPR-Cas9 technique in *A. fumigatus* (Figure S1). Morphological analysis in solid and liquid minimal media showed that the Δ*cnaA* mutant displayed defective morphological phenotypes including reduced hyphal growth and abnormal branching (Figure 1A). Notably, the abnormal hyphal phenotypes of the Δ*cnaA* mutant were exacerbated in the presence of 10 mM CaCl_2_, i.e., swollen hyphae tips, whereas no obvious differences were observed in the *cnaA^R^* complementary strain. Moreover, the Δ*cnaA* mutant showed a significant decrease in mycelial production compared to that of the wild-type at indicated time-points when cultured in media with or without 10 mM CaCl_2_ (Figure 1B). Interestingly, the calcineurin mutants ceased growth when incubated in the presence of calcium after 72 h, suggesting that the addition of calcium is toxic to the Δ*cnaA* mutant (Figure 1C). To confirm this toxicity, we cultured the wild-type and Δ*cnaA* strains in liquid minimal media (MM) with or without 10 mM CaCl_2_ for 96 h. Subsequently, the homogenized mycelia were inoculated into fresh liquid MM for another 48 h. The wild-type strains pretreated with or without calcium grew relatively well after transferring to fresh liquid MM. In contrast, the growth of the Δ*cnaA* mutant pretreated with calcium was dramatically decreased compared to that of the untreated Δ*cnaA* mutant (Figure 1D and 1E), confirming that the relatively low levels of extracellular calcium significantly inhibited the growth of the calcineurin mutant. Moreover, the Δ*cnaB* mutant exhibited identical phenotypes to those of the Δ*cnaA* mutant under the conditions we tested (Figure S2). Taken together, these data suggest that extracellular calcium is toxic to *A. fumigatus* calcineurin mutants and that calcineurin is essential for the survival of *A. fumigatus* in response to calcium stimuli.

**Figure 1.**
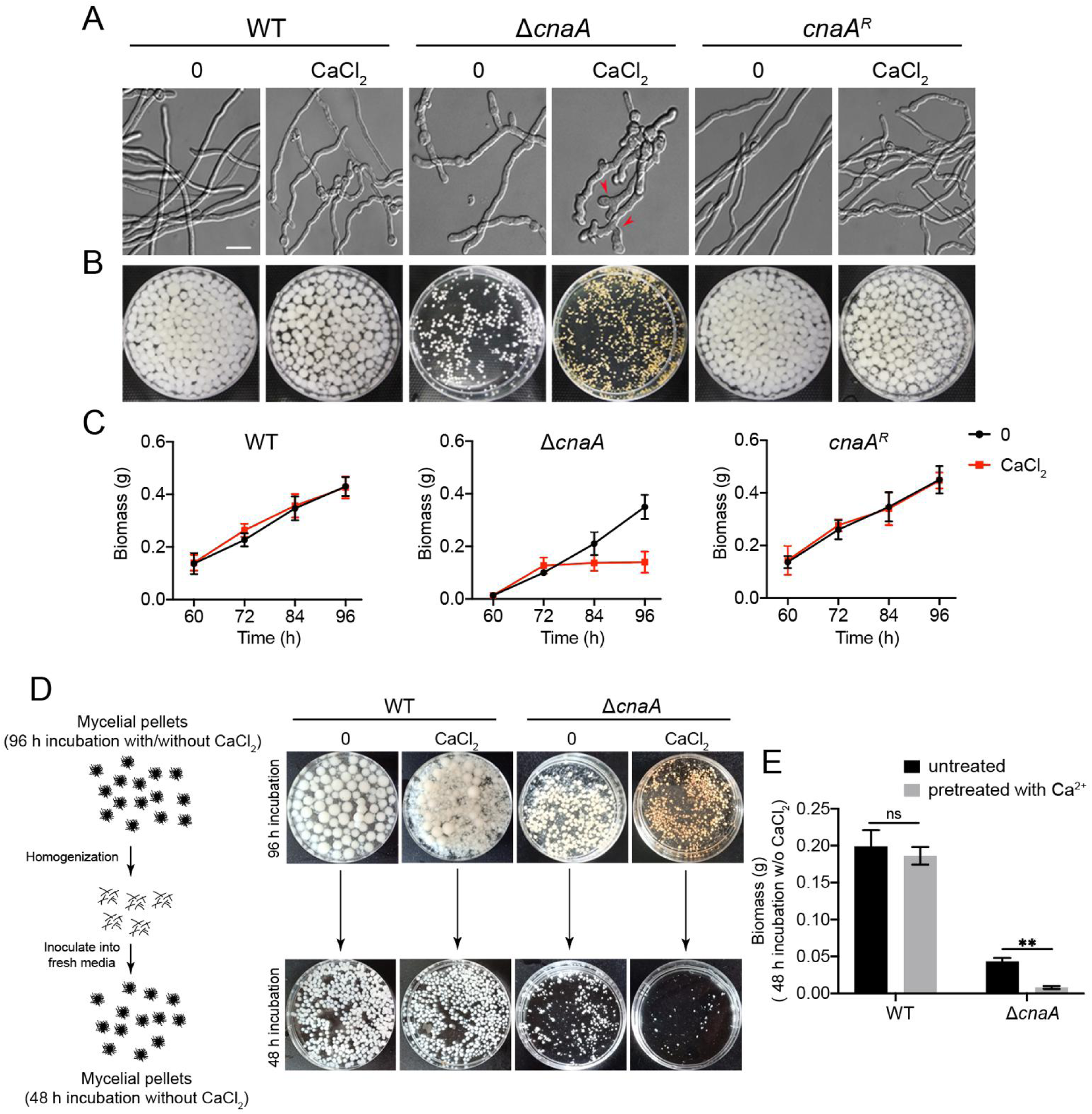
Calcium is toxic to calcineurin mutants. A. Differential interference contrast (DIC) images of hyphae grown in liquid MM with or without 10 mM CaCl_2_ at 37°C in stationary culture for 14 h. Scale bar represents 10 μm. B. The morphology of mycelial pellets of the indicated strains grown in MM with or without 10 mM CaCl_2_ at 37°C for 60 h. C. Quantification of biomass production for the wild-type, Δ*cnaA*, and *cnaA^R^* complementary strains grown in MM with or without 10 mM CaCl_2_ at different time points (60, 72, 84, and 96 h). D. Schematic diagram for determination of calcium toxicity to *A. fumigatus* cells. The wild-type and Δ*cnaA* strains were cultured in liquid MM with or without 10 mM CaCl_2_ at 37°C for 96 h. Then, the harvested mycelia were homogenized and inoculated into fresh liquid MM for another 48 h at 37°C. E. Quantification of biomass production for the wild-type and Δ*cnaA* strains pretreated with or without 10 mM CaCl_2_ grown in liquid MM at 37°C for 48 h. Statistical significance was determined by Student’s *t*-test. \*\**p* < 0.01.

### Δ*cnaA* mutant exhibits increased cytosolic and vacuolar calcium accumulation in response to extracellular calcium stimuli

Given that calcineurin is critical for regulating *A. fumigatus* survival in response to extracellular calcium, to further explore the mechanisms by which calcineurin regulates calcium homeostasis in *A. fumigatus*, we constructed wild-type and calcineurin mutants expressing codon-optimized aequorin to perform real-time monitoring of the dynamics of free Ca^2+^ concentration [Ca^2+^] in living hyphal cells (*Nelson et al., 2004*). Upon treatment with 10 mM CaCl_2_, the [Ca^2+^]_c_ (the free Ca^2+^ concentration in cytosol) in the wild-type strain transiently increased from a basal resting level of approximately 0.1 μM to a peak concentration of 0.63 μM and then gradually returned to a stable resting level (Figure 2A); thus, the amplitude of [Ca^2+^]_c_ between the peak and the resting status was 0.53 μM. In comparison, Δ*cnaA* showed a two-fold increased basal resting level of [Ca^2+^]_c_ (0.2 μM ± 0.04) and 26% increased amplitude of [Ca^2+^]_c_ (0.94 μM ± 0.21) under the same conditions, suggesting that cytosolic calcium homeostasis was affected by the deletion of *cnaA*. Next, we wondered whether the calcium equilibrium in mitochondria would be affected by calcineurin mutants. To test this possibility, we compared the [Ca^2+^]_mt_ (the free Ca^2+^ concentration in mitochondria) using a modified aequorin (Mt-Aeq) protein fused with a mitochondrial signal peptide. As shown in Figure 2B, the Δ*cnaA* mutant displayed a 20% increase in the basal resting level and a similar amplitude response of [Ca^2+^]_mt_ compared to the wild-type and *cnaA* complementing strains, suggesting that the deletion of *cnaA* only has a minor effect on mitochondrial calcium homeostasis. Next, we wondered in fungi, the vacuole might be a major intracellular Ca^2+^ store that plays a role in the regulation of Ca^2+^ homeostasis. However, until now, no reliable detection approach has been available for the real-time monitoring of the vacuolar free Ca^2+^ concentration ([Ca^2+^]_vac_) in fungi. To set up this system, we fused a guide sequence encoding the vacuolar lumen marker protein carboxypeptidase Y (CpyA) at the N-terminus of aequorin to generate the fusion protein CpyA-Aeq (*Kikuma, Ohneda, Arioka, & Kitamoto, 2006*). To verify the localization of the guided protein CpyA in *A. fumigatus*, the CpyA-GFP/RFP strain was generated as control. As shown in Figure 2C, the localization of the N-terminal CpyA-GFP fusion protein shows a vacuolar pattern. Further colocalization analysis showed that the previously verified vacuole membrane-localized protein, PmcA (the homolog of yeast P-type Ca^2+^-ATPase Pmc1), surrounded the CpyA-RFP (Figure 2C), demonstrating that CpyA acts as a typical vacuole lumen protein. Next, to further confirm the feasibility of the [Ca^2+^]_vac_ measurement system, we generated the overexpression strains of the vacuolar P-ATPases PmcA and PmcB, which are responsible for calcium influx into the vacuole by consuming ATP, and assessed the calcium levels in the vacuole. As shown in Figure S3, overexpressing *pmcA/B* remarkably elevated the basal resting level of [Ca^2+^]_vac_ and enhanced calcium influx into the vacuole compared with the wild-type strain, confirming the reliability of this vacuolar calcium measurement method. Thus, we employed the CpyA-Aeq system to monitor the free Ca^2+^ concentration of vacuoles. Interestingly, the basal resting [Ca^2+^] level in vacuoles (0.65 μM) prior to stimuli was significantly greater than that in the cytosol and mitochondria (Figure 2D), confirming that the vacuole is a major intracellular calcium store in *A. fumigatus*. Notably, in the Δ*cnaA* cells, the basal resting level of [Ca^2+^]_vac_ increased by 23% (0.8 μM), and the [Ca^2+^]_vac_ amplitude sharply increased to a 4.6-fold higher value (4.6 μM) compared to that of in wild-type cells upon exposure to 10 mM Ca^2+^ (Figure 2D). Collectively, these data suggested that the Δ*cnaA* mutant exhibits a dramatic increase in vacuolar Ca^2+^ influxes and induces excessive vacuolar Ca^2+^ accumulation.

**Figure 2.**
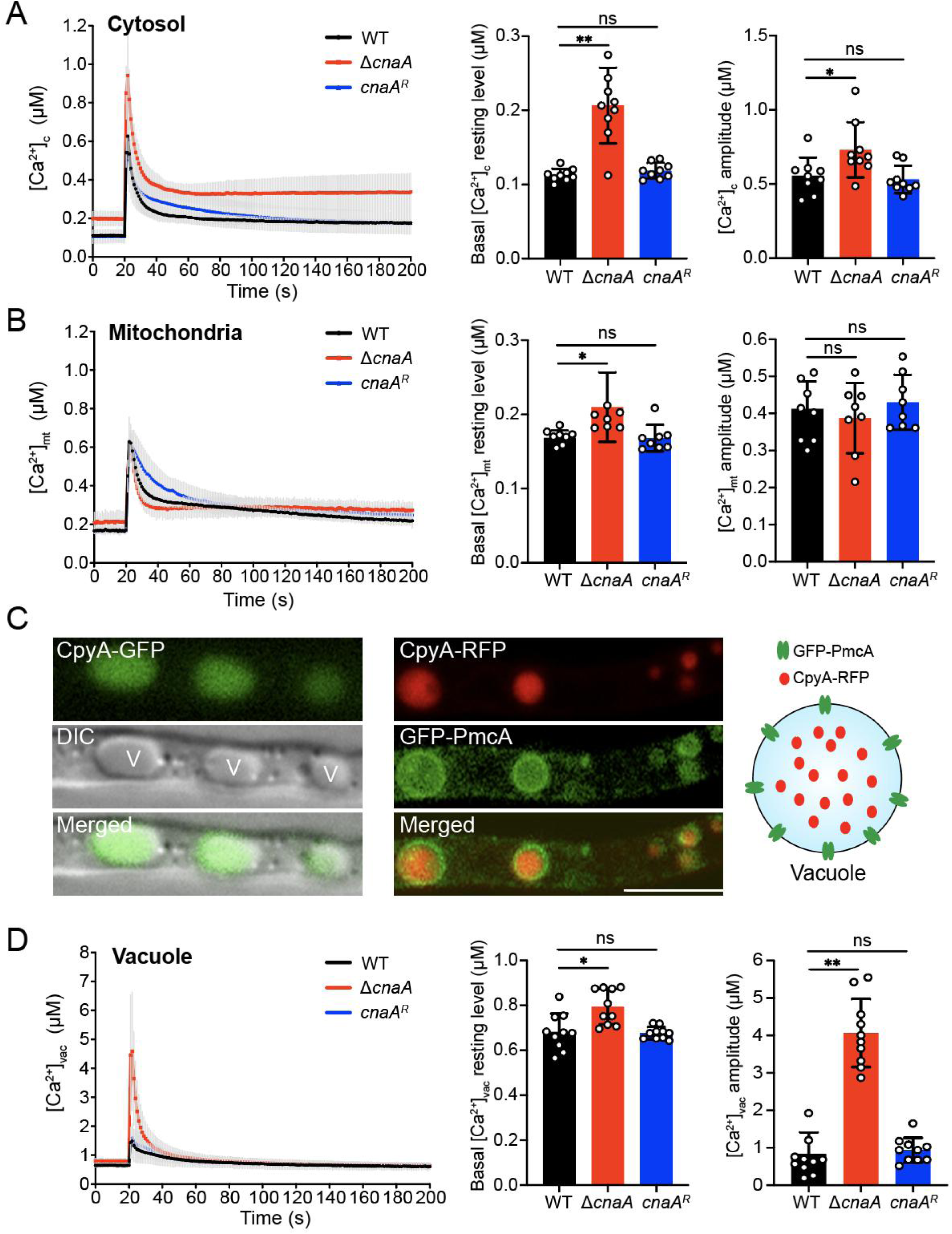
Deletion of *cnaA* affects intracellular calcium accumulation in response to extracellular calcium stimuli. A, B, and D. The linear graphs indicate the real-time [Ca^2+^]_c_, [Ca^2+^]_mt_, and [Ca^2+^]_vac_ changes in response to calcium stimuli. [Ca^2+^]_c_, the free Ca^2+^ concentration in cytosol; [Ca^2+^]_mt_, the free Ca^2+^ concentration in mitochondria; [Ca^2+^]_vac_, the free Ca^2+^ concentration in vacuoles. Basal [Ca^2+^], the resting level prior to extracellular calcium stimulus, and [Ca^2+^] amplitude, the difference between the basal [Ca^2+^] level and the poststimulatory peak value, are presented. Data are the average of at least six experiments. Error bars show the standard deviation. Statistical significance was determined by Student’s *t*-test. \**p* < 0.05; \*\**p*< 0.01; ns, not significant. C. The localization of CpyA. The letter V in the DIC image indicates vacuole (Left). The localization analysis (Middle) and schematic illustration (Right) of PmcA and CpyA. Scale bar represents 5 μm.

### Extracellular calcium induces vacuolar fragmentation and autophagy in the Δ*cnaA* mutant

Next, we wondered whether the excessive vacuolar Ca^2+^ accumulation in the Δ*cnaA* mutant would cause any changes of vacuolar morphology. As shown in Figure 3A, fluorescence microscopy analysis showed that wild-type and Δ*cnaA* cells had similar oval-shaped mature vacuoles as indicated by CpyA-GFP when cultured in liquid minimal media for 36 h with or without the addition of calcium. Notably, when incubation time was prolonged to 60 h, the Δ*cnaA* mutant showed highly fragmented fluorescence patterns of vacuoles in the presence of 2.5 or 10 mM calcium (Figure 3A). In comparison, in the absence of calcium treatment, the vacuolar morphologies of Δ*cnaA* mutants showed no detectable differences compared to the parental strain. These data indicate that significant vacuolar fragmentation occurs in the Δ*cnaA* mutant when grown in liquid minimal media containing extra calcium. Because the vacuole is the major site for macromolecular degradation through autophagy, to examine whether the deletion of *cnaA* would trigger autophagy under extracellular calcium condition, we generated double-labeled strains with the autophagy marker Atg8 tagged at its N-terminus with GFP and the vacuole marker CpyA fused with RFP and monitored these in the Δ*cnaA* and wild-type backgrounds. As shown in Figure 3B, GFP-Atg8 fusion proteins show a punctate pattern that is proximal to the CpyA-RFP-labeled vacuoles in the wild-type and Δ*cnaA* strains when incubated for 36 h in the presence or absence of calcium. However, when cultures were stimulated by the addition of CaCl_2_ for 60 h, the green fluorescence patterns of GFP-Atg8 overlapped with the CpyA-RFP-labeled vacuoles (Figure 3B). Furthermore, Western blotting showed that the Δ*cnaA* strain displayed increased cleavage of GFP from the GFP-Atg8 fusion protein compared to the wild-type cells when cultured for 60 h under the control of the constitutive *gpdA* promoter (Figure 3C) or the native *atg8* promoter (Figure S4A-S4B) in the presence of calcium. In addition, cleavage of GFP was not significantly exacerbated when media were supplemented with other divalent metal cations including magnesium, zinc, iron, and copper (Figure S4C), suggesting that the autophagy was specifically induced by calcium. Next, to examine the contribution of autophagy to Ca^2+^ toxicity in the Δ*cnaA* mutant, we deleted the classic autophagy-dependent gene *atg2*, which encodes a peripheral membrane protein for autophagic vesicle formation (*Kotani, Kirisako, Koizumi, Ohsumi, & Nakatogawa, 2018*; *Tang, Takahashi, & Wang, 2019*), in the Δ*cnaA* background. As shown in Figure 3D, GFP-Atg8 consistently exhibited a punctate pattern and did not co-localize with the vacuolar marker CpyA-RFP in the Δ*cnaA*Δ*atg2* mutant. Furthermore, Western blotting showed that no (or an undetectable level of) GFP was cleaved from GFP-Atg8 fusion protein in the Δ*cnaA*Δ*atg2* double mutant compared with the Δ*cnaA* mutant (Figure 3E), suggesting that autophagy was suppressed in the Δ*cnaA*Δ*atg2* mutant. Notably, as shown in Figure 3F, the Δ*cnaA*Δ*atg2* mutant displayed a delay in growth cessation (at 84 h) compared to that of the Δ*cnaA* mutant (at 72 h) under calcium treatment conditions (Figure 3F). However, the Δ*cnaA*Δ*atg2* mutant pretreated with or without calcium showed a comparable biomass production compared to that of the Δ*cnaA* mutant after homogenization and transfer to fresh liquid MM, suggesting that the deletion of *atg2* did not prevent the cell death induced by a lack of *cnaA* after culture (Figure 3G), indicating that Atg2 is required for autophagy but not Ca^2+^ toxicity in the calcineurin-deficient mutant upon calcium stimuli. Taken together, these data suggest that a lack of CnaA results in specific calcium-induced autophagy in *A. fumigatus*.

**Figure 3.**
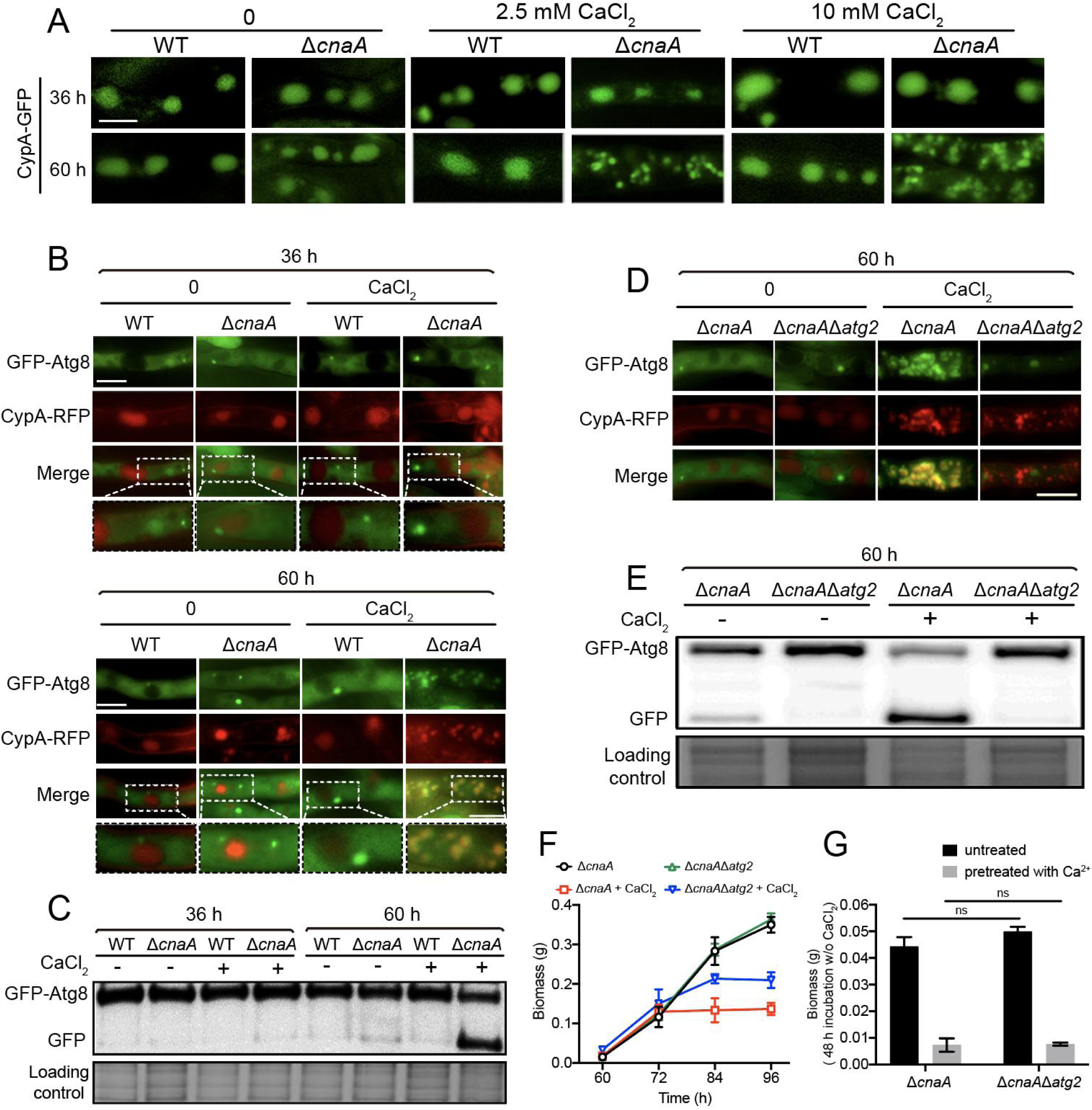
The loss of CnaA causes vacuolar fragmentation and *atg2*-related autophagy in the presence of excess calcium. A. Vacuolar morphology was monitored by fluorescence microscopy using CpyA-GFP as vacuolar lumen marker. The indicated strains were grown in liquid MM with or without calcium CaCl_2_ for 36 and 60 h. Scale bar represents 5 μm. B and D. The localization of CpyA-RFP and GFP-Atg8 was used to monitor the autophagy process. When autophagy is induced, the outer membrane of the autophagosome was fused with the membrane vacuole, whereas the inner membranes and the cargo were delivered into the vacuolar lumen for degradation. In this process, the Atg8 protein transferred from the autophagosome to the vacuole and was cleaved. The GFP/RFP signals in the mycelium pellets were observed after the indicated strains were cultured in a rotary shaker for 36 or 60 h. Scale bars represent 5 μm. C and E. Western blotting showed the GFP-Atg8 cleavage of the related strains in MM with or without 10 mM CaCl_2_ for 36 or 60 h. The extent of autophagy was estimated by calculating the amount of free GFP compared to the total amount of GFP-Atg8 and free GFP. Coomassie blue staining served as a loading control. F. Quantification of biomass production for the Δ*cnaA* and Δ*cnaA*Δ*atg2* strains grown in MM with or without 10 mM CaCl_2_ at different time points (60, 72, 84, and 96 h). G. Quantification of biomass production for the Δ*cnaA* and Δ*cnaA*Δ*atg2* strains pretreated with or without 10 mM CaCl_2_ grown in liquid MM at 37°C for 48 h. Statistical significance was determined by Student’s *t*-test. ns, not significant.

### Lack of the calcium channel CchA alleviates the calcium toxicity-related phenotypes in the *cnaA* null mutant

Our previous studies in a model fungus *A. nidulans* indicated that calcineurin negatively regulates CchA (the voltage-gated Ca^2+^ channel component of high-affinity calcium influx system) upon calcium uptake in response to extracellular calcium; the deletion of *cchA* suppressed the hypersensitivity to external calcium in calcineurin mutants (S. *Wang, Liu, Qian, Zhang, & Lu, 2016*). We hypothesized that a similar situation would occur in *A. fumigatus*. As expected, *cchA* deletion partially suppressed the colony radial growth defects and hyperbranching polarity growths in *cnaA* deletion mutants with or without calcium treatment (Figure S5). Notably, a lack of CchA significantly rescued the defective hyphal growth of the *cnaA* mutant in calcium treatment conditions (Figure 4A and 4B). The detection of calcium dynamics in living cells showed that the [Ca^2+^]_vac_ amplitude and the basal resting [Ca^2+^]_c_ of the Δ*cnaA*Δ*cchA* mutant were approximately 53% and 20% lower, respectively, than those in the Δ*cnaA* mutant (Figure 4C and 4D), suggesting that calcineurin mediates cytosolic and vacuolar calcium homeostasis by negatively regulating CchA. Accordingly, the colocalization of GFP-Atg8 and CpyA-RFP (vacuolar marker) and the vacuolar fragmentation were not observed in the Δ*cnaA*Δ*cchA* mutant (Figure 4E). Moreover, Western blots showed that the Δ*cnaA*Δ*cchA* mutant exhibited an autophagy level similar to that of the wild-type strain (Figure 4F), suggesting that autophagy in response to external calcium is inhibited when *cchA* is deleted in the Δ*cnaA* mutant. These data suggest that a lack of the calcium channel CchA alleviates the calcium toxicity-related phenotypes in the *cnaA* null mutant.

**Figure 4.**
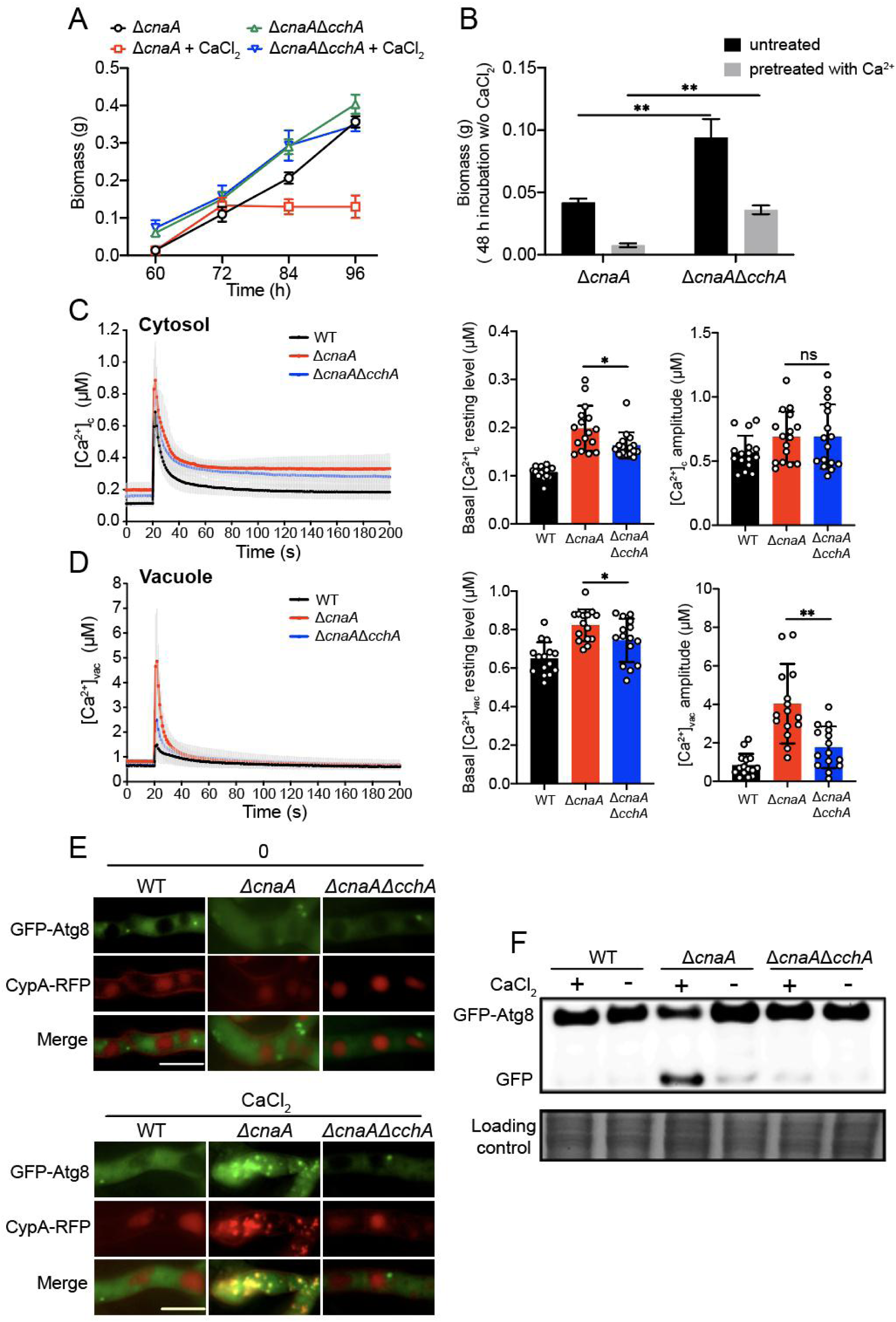
Deletion of *cchA* alleviates the calcium toxicity-related phenotypes in the Δ*cnaA* mutant. A. Quantification of biomass production for the Δ*cnaA* and Δ*cnaA*Δ*cchA* strains grown in MM with or without 10 mM CaCl_2_ at different time points (60, 72, 84 and 96 h). B. Quantification of biomass production for the Δ*cnaA* and Δ*cnaA*Δ*cchA* strains pretreated with or without 10 mM CaCl_2_ grown in liquid MM at 37°C for 48 h. Statistical significance was determined by Student’s *t*-test. \*\**p* < 0.01. C and D. The comparison of [Ca^2+^]_c_ and [Ca^2+^]_vac_ of WT, Δ*cnaA* and Δ*cnaA*Δ*cchA* in resting and dynamic level. \**p* < 0.05; \*\**p* < 0.01; ns, not significant. E and F. The co-localization analysis of CpyA-RFP and GFP-Atg8 and Western blotting showed the localization and cleavage of GFP-Atg8 of the related strains in MM with or without 10 mM CaCl_2_ for 60 h. Scale bars represent 5 μm.

### Lack of *CchA* partially restores the virulence of the Δ*cnaA* mutant in a *Galleria mellonella* infection model but not in a mouse infection model

The aforementioned data demonstrated that the absence of CchA in the Δ*cnaA* mutant partially suppressed the growth defects of *A. fumigatus in vitro*. Next, we further tested whether the Δ*cnaA*Δ*cchA* mutant would affect virulence in *in vivo* animal models. As shown in Figure 5A, in the insect infection model of the wax moth *Galleria mellonella*, the Δ*cnaA*Δ*cchA* mutant had a significantly lower survival rate (40%) than that of the Δ*cnaA* mutant (90%), suggesting that a lack of *cchA* partially restored the virulence of the Δ*cnaA* mutant. Furthermore, the virulence of the Δ*cnaA*Δ*cchA* mutant was tested in a mouse model. Immunocompromised ICR mice were infected with fresh conidia, homogenized hyphae, or luciferase-probed related strains by an intratracheal route. After two weeks, the mice infected with wild-type *A. fumigatus* display high mortality, whereas both the Δ*cnaA*Δ*cchA* and Δ*cnaA* mutants caused low mortality (Figure 5A). Statistical analysis showed that there was no significant difference between Δ*cnaA*Δ*cchA* and Δ*cnaA* in survival rates for either the conidia or the homogenized hyphal infection model. Histopathological examinations and live animal imaging showed that the lungs from mice inoculated with the wild-type fungus displayed aggressive fungal growth and luminescence signals around the lungs (Figure 5B and 5C). In contrast, there was no detectable fungal growth or luminescence signals in the lungs of mice infected with the Δ*cnaA*Δ*cchA* or Δ*cnaA* mutants, implying that the immune system of mice might be able to eliminate inoculated conidia or hyphae. These data suggest that a lack of CnaA results in the loss of virulence in immunocompromised mice, and the deletion of *cchA* partially restored virulence in wax moths but not in the mouse model.

**Figure 5.**
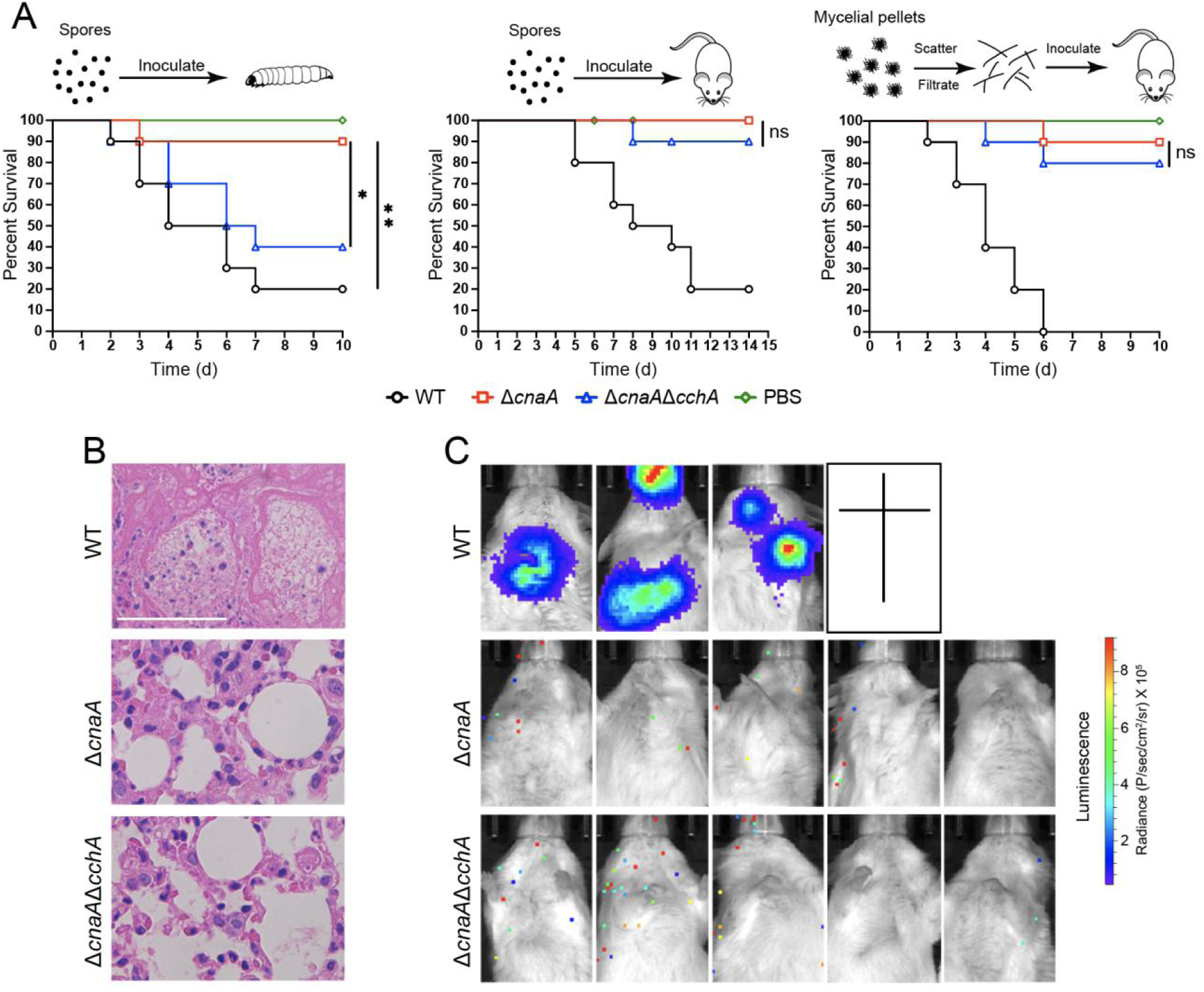
Deletion of *cchA* in the Δ*cnaA* mutant partially restores the virulence in a *Galleria mellonella* infection model but not in a mouse infection model. A. Survival curves of *G. mellonella larvae* infected with WT, Δ*cnaA*, and Δ*cnaA*Δ*cchA*. Larvae were infected with 8 x 10^5^ conidia, and PBS-injected larvae served as controls (left panel). Survival curve of mice infected with conidia or hyphae (middle and right panels) of WT, Δ*cnaA*, and Δ*cnaA*Δ*cchA A. fumigatus*. Statistical analysis between groups were using the log-rank test. \**p* < 0.05; \*\**p* < 0.01; ns, not significant. B. Histopathological sections of lung tissue from sacrificed mice infected with conidia of each strain. Periodic acid-Schiff stain was used to visualize fungal growth. Scale bar represents 50 μm. C. Luciferase signals in mice infected with hyphae were measured with a live animal imaging system 4 days after infection.

### Overexpressing P-type Ca^2+^-ATPase PmcA significantly rescued the calcium toxicity-related phenotypes in the Δ*cnaA* mutant

In fungi, the vacuole serves as the major Ca^2+^ store and plays a critical role in the detoxification of cytoplasmic Ca^2+^. Vacuolar calcium equilibrium is maintained by the P-type Ca^2+^-ATPase Pmc family and the H^+^/Ca^2+^ exchanger Vcx family, which are regulated by calcineurin (*Cyert & Philpott, 2013*). In line with this information, the Δ*cnaA* mutant exhibited lower expression levels of *pmcA* and *pmcC* compared to those of the wild-type, whereas the transcript level of vacuolar H^+^/Ca^2+^ exchanger family members, especially *vcxB*, *vcxD*, and *vcxE*, was upregulated in *A. fumigatus* (Figure S6), demonstrating that calcineurin positively regulates the P-type Ca^2+^-ATPase Pmc family while negatively regulating the H^+^/Ca^2+^ exchanger Vcx family. Given the calcium toxicity-related phenotypes observed in the Δ*cnaA* mutant, we hypothesized that the upregulation of H^+^/Ca^2+^ exchanger Vcx family members in the Δ*cnaA* mutant would be insufficient to detoxify excess cytoplasmic calcium in response to external calcium, and thus a Δ*cnaA^OE::pmcA^* strain overexpressing PmcA, the yeast P-type Ca^2+^-ATPase Pmc1 homolog in *A. fumigatus*, under the control of the strong constitutive promoter P_gpdA_ was used in the Δ*cnaA* background to test whether the defective phenotypes in the Δ*cnaA* mutant would be suppressed. qRT-PCR analysis confirmed that *pmcA* was highly overexpressed in the Δ*cnaA* mutant (Figure S3). Further live-cell detection of [Ca^2+^]_vac_ showed that the Δ*cnaA^OE::pmcA^* strain displayed an elevated basal resting level of [Ca^2+^]_vac_ and had an enhanced [Ca^2+^]_vac_ amplitude compared to that of the Δ*cnaA* mutant (Figure 6A), indicating that calcium was effectively transported from the cytoplasm to vacuoles by PmcA. Additionally, autophagy assays showed that the Δ*cnaA^OE::pmcA^* strain displayed reduced autophagy compared to that of the Δ*cnaA* mutant in response to external calcium stimuli (Figure 6B and 6C). Moreover, the overexpression of *pmcA* in the *cnaA* mutant markedly suppressed the growth cessation in calcium treatment conditions (Figure 6D). The Δ*cnaA^OE::pmcA^* strain pretreated with or without calcium (96 h) showed dramatically increased biomass production compared to that of the Δ*cnaA* mutant after homogenization and transfer to fresh liquid MM (Figure 6E). Thus, these data suggested that overexpressing P-type Ca^2+^-ATPase PmcA partially rescues calcium toxicity-related phenotypes in the Δ*cnaA* mutant.

**Figure 6.**
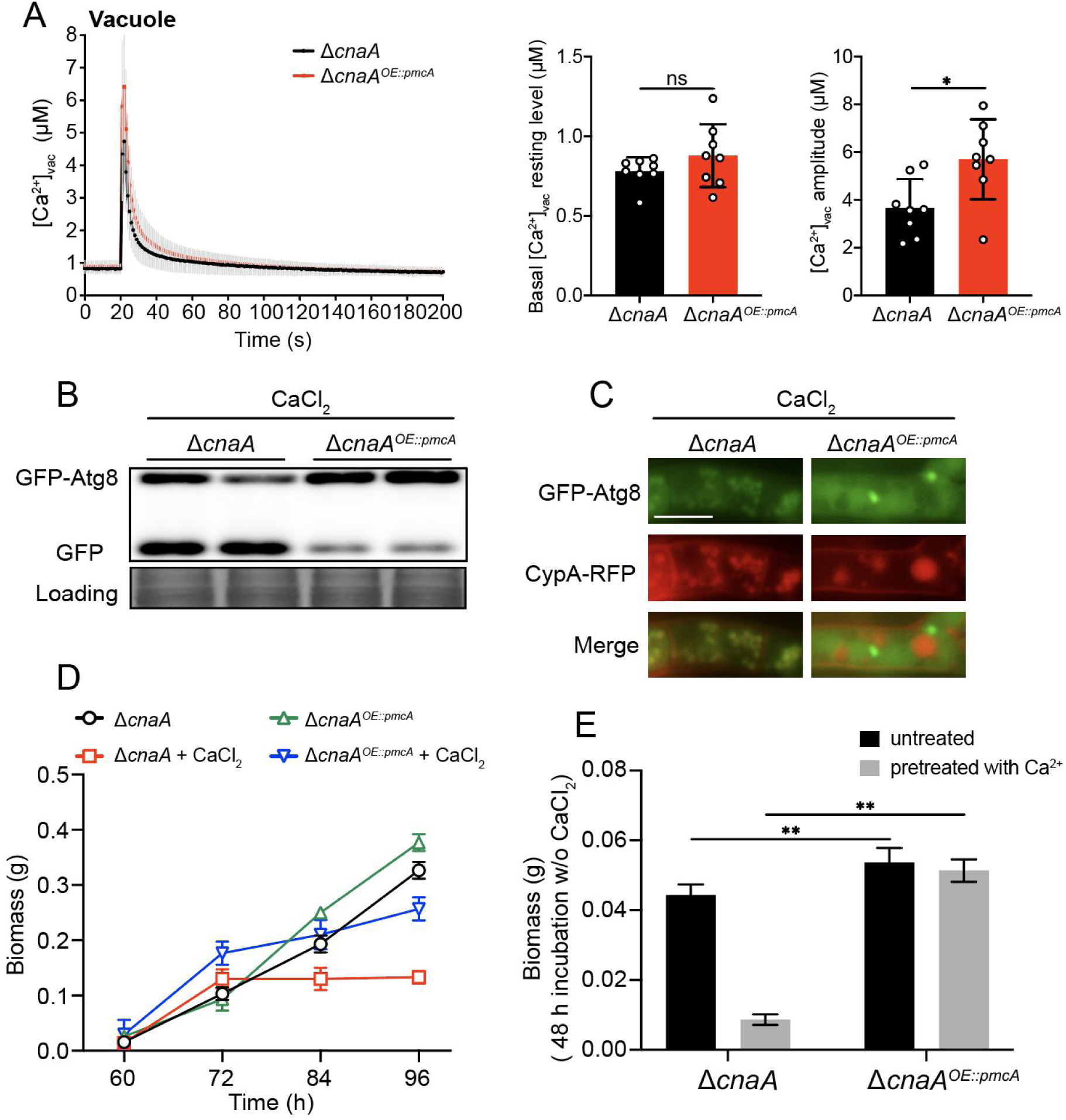
Overexpressing P-type Ca^2+^-ATPase PmcA in the Δ*cnaA* mutant significantly rescues calcium toxicity-related phenotypes. A. The comparison of [Ca^2+^]_vac_ of Δ*cnaA* and Δ*cnaA*^OE::pmcA^ in resting and dynamic level. B and C. Western blots analysis and co-localization analysis of CpyA-RFP and GFP-Atg8 were used to estimate the autophagy level in Δ*cnaA* and Δ*cnaA^OE::pmcA^* strains. Scale bar represents 5 μm. D. Quantification of biomass production for the Δ*cnaA* and Δ*cnaA^OE::pmcA^* strains grown in the indicated conditions at different time points (60, 72, 84 and 96 h). E. Quantification of biomass production for the Δ*cnaA* and Δ*cnaA^OE::pmcA^* strains pretreated with or without 10 mM CaCl_2_ grown in liquid MM at 37°C for 48 h. Statistical significance was determined by Student’s *t*-test. \*\**p* < 0.01.

### The Δ*cnaA* mutant exhibits global fragmentation of nuclei and organelles upon calcium stimuli

To further dissect the molecular mechanisms operating in the *A. fumigatus* Δ*cnaA* mutant upon *in vitro* calcium stimuli, we performed proteomic comparison analysis using tandem mass tags (TMTs) and liquid chromatography tandem MS (LC/MS-MS) and compared wild-type and Δ*cnaA* strains grown in the presence of extracellular calcium. Proteins with change ratios significantly different from general protein variation (1.50 < Δ*cnaA*/wild-type < 0.50) were selected for gene ontology (GO) annotation and enrichment analysis. As shown in Table S3, the expression levels of 618 and 218 proteins were increased and decreased, respectively, in the Δ*cnaA* mutant compared to the wild-type strain. Most upregulated proteins were involved in metabolic processes of biomolecules, especially in catabolism and proteolysis (Figure S7A). In contrast, most of the downregulated proteins were associated with biosynthesis and/or the transport of biomolecules and inorganic ions (Figure S7B). Kyoto Encyclopedia of Genes and Genomes (KEGG) annotation revealed that the upregulated proteins were primarily associated with various amino acid metabolism pathways, especially tyrosine, beta-alanine, valine, and leucine (Figure 7A), whereas the downregulated proteins mainly focused on biosynthesis-related pathways including ribosome (protein synthesis), thiamine metabolism, and steroid biosynthesis. Of note, the upregulated enriched group in the Δ*cnaA* mutant contained 23 peptidases, 18 hydrolases, 4 nucleases, 3 proteases, 3 autophagy-related proteins, and 1 metacaspase (Figure 7A), which may lead to global protein degradation and cause apoptosis/apoptotic-like programmed and/or autophagic cell death in Δ*cnaA* cells under calcium treatment. Thus, we hypothesized that CnaA-involved cellular calcium equilibrium may regulate whole-cell organelle networks. To further assess whether a lack of CnaA would induce abnormal organelle and nuclear morphology with calcium treatment, we probed the nucleus (RFP-H2A), endoplasmic reticulum (Erg11A-GFP), Golgi apparatus (RFP-PH^OSBP^), and mitochondria (MrsA-RFP). As predicted, fluorescence observation showed that these organelles displayed fragmentation that was comparable to that of the vacuole in the Δ*cnaA* mutant after calcium treatment for 60 h (Figure 7C); apoptosis/apoptotic-like and/or autophagic cell death may occur. A similar phenomenon was observed in *A. fumigatus* cells treated with the calcineurin inhibitor FK506 (Figure 7D). Taken together, these data suggested that global nuclear and organelle fragmentation may contribute to calcium toxicity in mutants lacking functional calcineurin.

**Figure 7.**
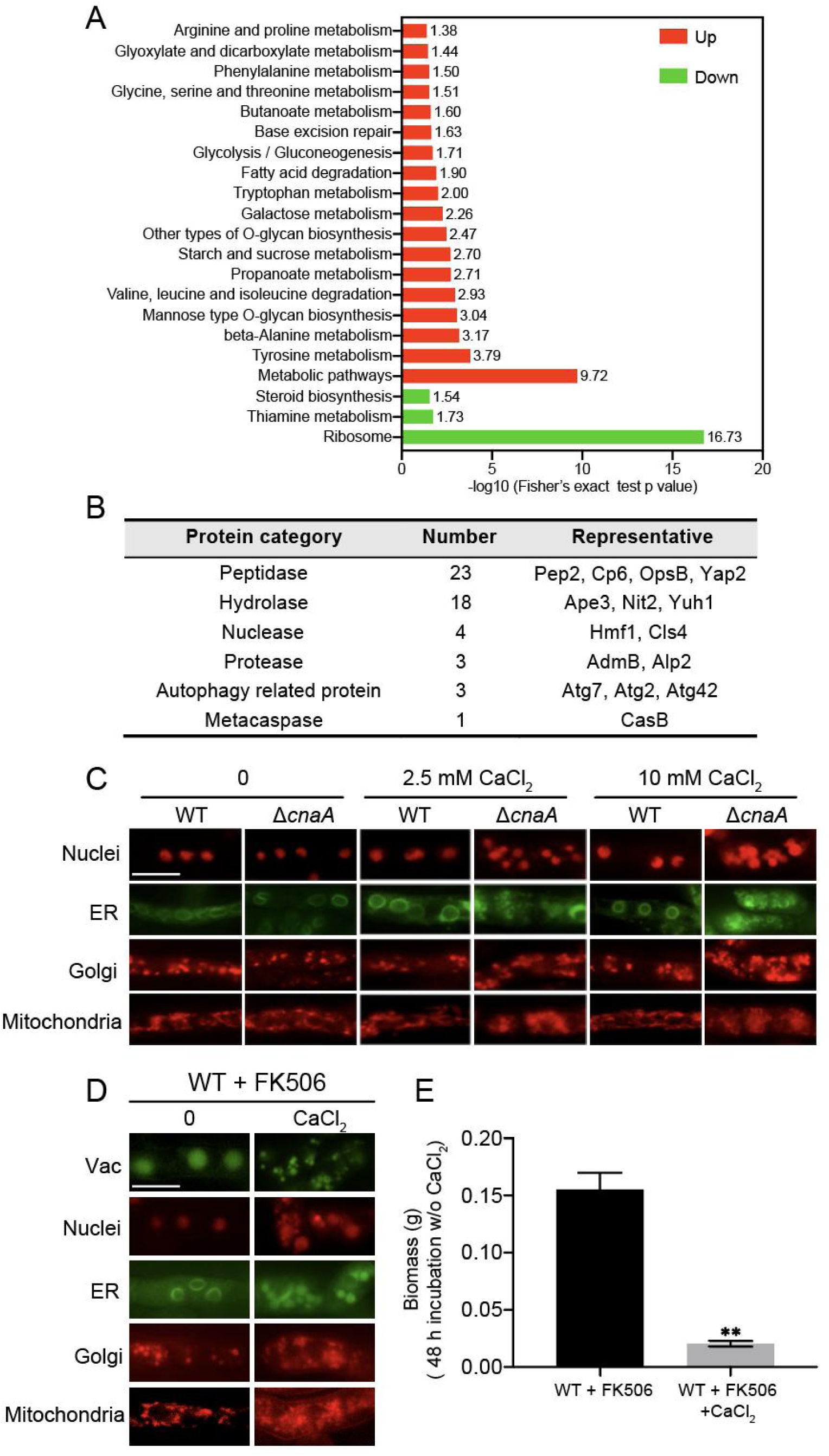
Δ*cnaA* mutant shows global fragmentation of nuclei and organelles upon calcium stimuli. A. KEGG pathways enriched in proteins with more than 1.5-fold changes between WT and Δ*cnaA*. Red and green items indicate upregulated and downregulated pathways, respectively. The KEGG online service tool KAAS (http://www.genome.jp/kaas-bin/kaas_main) was used to annotate protein KEGG database descriptions. B. Selected enzymes related to material degradation in the upregulated set of proteins are presented. C. The organelle morphologies of the wild-type and Δ*cnaA* strains grown in liquid MM with or without calcium for 60 h. Scale bar represents 5 μm. D. The organelle morphologies of the wild-type strain grown in liquid MM supplemented with FK506 in the presence or absence of 2.5 mM CaCl_2_ for 60 h. A 50 μL sample of 2 mM FK506 was added into 100 ml liquid MM at 0 and 48 h during culture. Scale bar represents 5 μm. E. Quantification of biomass production for the wild-type strain pretreated with or without 10 mM CaCl_2_ grown in liquid MM supplemented with FK506 at 37°C for 48 h. Statistical significance was determined by Student’s *t*-test. \*\**p* < 0.01.

## Discussion

The critical role of calcineurin in calcium dysregulation-related human diseases has been recognized for decades (*Y. J. Park, Yoo, Kim, & Kim, 2020*). In fungi, calcineurin has been considered an attractive antifungal target because it is essential for stress survival, development, drug resistance, and virulence (*Juvvadi et al., 2017*). Previous studies have reported that calcineurin protects *Candida albicans* from toxicity caused by the endogenous levels of calcium present in serum (*Blankenship & Heitman, 2005)*. However, the underlying mechanism by which calcineurin orchestrates intracellular calcium dynamics remains elusive. Through the successful development of an intracellular calcium dynamics monitoring method in this study, we demonstrated that calcineurin dysfunction disturbed calcium homeostasis in the cytosol, vacuole, and mitochondria, leading to a dramatic induction of autophagy and organelle fragmentation and ultimately resulting in significant cell growth inhibition and cell death in the presence of calcium stimuli.

Typically, the resting cytosolic Ca^2+^ concentration in *A. fumigatus* ranges from 0.05 to 0.1 μM. In the Δ*cnaA* mutant, the cytosolic resting Ca^2+^ level increased by two-fold compared to that of the wild-type prior to calcium stimuli, which is in line with our previous reports on *A. nidulans*. Moreover, when cells were challenged with 10 mM Ca^2+^, the Δ*cnaA* mutant displayed a 33% increase in [Ca^2+^]_c_ amplitude compared to that of the wild-type. These data support the evolutionarily conserved role of calcineurin in maintaining cytosolic calcium homeostasis across fungal species. The sustained increased cytosolic calcium concentration led us to question whether calcium homeostasis in the intracellular calcium stores such as mitochondria and vacuole would be affected by a lack of calcineurin. We established a novel vacuolar Ca^2+^ detection approach by fusing the vacuole-targeted protein carboxypeptidase Y (CpyA) to the N-terminus of aequorin, and we found that in the presence of extracellular calcium stimuli, the basal resting level of [Ca^2+^]_vac_ and the transient [Ca^2+^]_vac_ in the Δ*cnaA* mutant were significantly increased, whereas the calcium signature in mitochondria was not significantly affected, implying that vacuoles may be the major site for the detoxification of excess cytosolic calcium in *A. fumigatus*. Collectively, these data demonstrated that calcineurin not only regulates the resting cytosolic calcium but also affects the vacuolar calcium transient response. Importantly, the increased cytosolic resting [Ca^2+^]_c_ level and the [Ca^2+^]_vac_ amplitude seen in the Δ*cnaA* mutant can be suppressed by deleting the calcium channel CchA, suggesting that calcineurin negatively regulates CchA when cells are exposed to calcium. Accordingly, the feedback relationship between calcineurin and the calcium channel CchA has been described in yeast and *A. nidulans* (*Ma et al., 2011;* S. *Wang et al., 2016*). In yeast, it has been proposed that Cch1 is dephosphorylated by calcineurin *in vitro* (*Bonilla & Cunningham, 2003*), leading to the inhibition of calcium channel activity, which is in accord with our phosphoproteomic analysis showing that Serine 55 in CchA could be directly dephosphorylated by calcineurin in *A. fumigatus* (Fig. S8).

In addition to the disordered intracellular calcium homeostasis, the Δ*cnaA* mutant exhibited the global fragmentation of organelles including the vacuole, the nucleus, ER, and mitochondria, and aggressive autophagy was observed when the Δ*cnaA* mutant was grown in the presence of 10 mM calcium. This autophagy resulted in the cessation of growth and cell death with prolonged incubation time, indicating that the relatively low concentration of calcium is toxic to calcineurin-deficient mutant *A. fumigatus*. It has been well established that the vacuole plays a key role in detoxifying the excess calcium when cells are exposed to external calcium. Thus, we hypothesized that the cytoplasmic calcium excess combined with deficient vacuolar detoxification capacity would give rise to the calcium-toxicity related phenotypes observed in the Δ*cnaA* mutant. This hypothesis was supported by the fact that alleviating cytosolic calcium overload by the deletion of *cchA* or increasing vacuolar calcium storage capacity by overexpressing of the P-type Ca^2+^-ATPase PmcA can significantly suppress the defective phenotypes of the Δ*cnaA* mutant. These results highlighted the important roles of calcineurin in mediating calcium homeostasis for fungal survival and the detoxification functions of the vacuoles. Of note, blocking CchA or the overexpression of PmcA is unable to completely suppress the vacuolar fragmentation, aggressive autophagy, and growth cessation in the Δ*cnaA* mutant in the presence of calcium stimuli, implying that there are other calcium pumps or transporters may also be co-operated with them and involved in the regulation of calcium homeostasis. Probably, examples include the mitochondrial Ca^2+^ uniporter McuA, the Golgi-localized P-type Ca^2+^/Mn^2+^-ATPase PmrA, and the ER-localized P-type ATPase SpfA (*Pinchai et al., 2010*; *Song, Liu, Zhai, Huang, & Lu, 2016*; *Yu et al., 2012*), all of which have been demonstrated to be required for mediating calcium homeostasis in response to environmental stresses. In addition, although our results suggested that autophagy triggered by the dysregulation of intracellular calcium homeostasis in the Δ*cnaA* mutant correlates with cell death because blocking autophagy by *atg2* deletion could slow the cell death progress, we cannot eliminate the possibility that other “toxic factors” affect the viability of *A. fumigatus* calcineurin mutants in the presence of calcium. Proteomic analysis revealed that various peptidases, nucleases, metacaspases, and autophagy-related proteins were upregulated in the Δ*cnaA* mutant compared to the wild-type strain in calcium-treated conditions, implying that apoptosis/apoptotic-like and/or autophagic cell death may occur.

Although cyclosporine A and FK506 (tacrolimus), calcineurin inhibitors, have been shown to possess *in vitro* antifungal activity against fungal pathogens including *A. fumigatus*, *C. albicans*, and *C. neoformans*, the clinical use of calcineurin inhibitors for antifungal therapy remains limited due to their immunosuppressive side effects (*Fox & Heitman, 2002*; *Groll, De Lucca, & Walsh, 1998*; *Hamamoto et al., 2018*). Because calcineurin inhibitors exhibit synergistic fungicidal activity with antifungal drugs, research on analogues of calcineurin inhibitors with antifungal activity but low immunosuppressive properties may be a promising direction. Encouragingly, APX879, the FK506 analogue, has been shown to exhibit reduced immunosuppressive activity while maintaining limited antifungal activity against pathogens in a murine infection model (*Juvvadi et al., 2019*). Our findings showing that a relatively low concentration of calcium is sufficient to significantly inhibit the growth of calcineurin mutants highlights a possibility that the combination of calcium and nonimmunosuppressive calcineurin inhibitors could enhance therapeutic efficacy against human pathogens, and the addition of calcium may reduce the dosage of immunosuppressive agents needed, thus minimizing side effects.

## Materials and Methods

### Strains, media, and culture conditions

was listed in the supplementary data Table S1. Strains were routinely grown in the following media: minimal media (MM) containing 1% glucose, 2% agar, 1 ml/L trace elements and 50 ml/L 20 × salt solution, as described previously (*Zhang & Lu, 2017*). The pH was adjusted to 6.0 : MM supplemented with 5 mM uridine and 10 mM uracil were used for uracil and uridine auxotroph strains. The recipe for liquid glucose minimal media is identical to that for MM, except without agar. All strains were cultured at 37°C.

### Quantification of biomass production

10^5^ conidia were incubated in 100 ml of MM with or without 10 mM CaCl_2_ at 37°C in a rotary shaker at 200 rpm for 60, 72, 84, and 96 h. Biomass curves were drawn after mycelial pellets were collected, dried, and weighed.

### Assessment of hyphal survival

The related strains were grown in liquid MM with or without 10 mM CaCl_2_ with 220 rpm shaking at 37°C for 96 h. Subsequently, the mycelium pellets of each strain were crushed and homogenized into hyphae fragments using tissue grinding and filtered with lens-cleaning paper in sterile ddH_2_O. The optical density at 600 nm (OD600) of each group was finally adjusted to 0.2. Then, 100 μL of suspension containing hyphae fragments was inoculated into liquid MM for 48 h via shake cultivation, and the biomass was weighed and compared.

### Constructions for plasmids

For the construction of complementary *cnaA/cnaB,* p-zero-hph-cnaA plasmid was generated as follows: the selectable marker *hph* from PAN7-1 was amplified by PCR using the primers hph-F and hph-R and then cloned into the pEASY-Blunt vector (TransGen Biotech), generating the plasmid p-zero-hph. The primers cnaA/cnaB-notI-F and cnaA/cnaB-notI-R were used to generate a fragment that includes the promoter sequence, the complete ORF, and the 3’UTR of *cnaA/cnaB*. This fragment was then cloned into the *Not*I site of p-zero-hph to generate the plasmid p-zero-hph-cnaA/cnaB.

For measuring the calcium level in the cytosol and mitochondria, the plasmids pAMA1-P_gpdA_-Aeq and pAMA1-P_gpdA_-mt-Aeq were generated as follows: PCR was performed using the primers Ama1-BamHI-gpd-F and Ama1-BamHI-trpC-R to amplify P_gpdA_-Aeq-T_trp_ and P_gpdA_-mt-Aeq-T_trpC_ fragments from the existing plasmids pAEQS1-15 and pAEQ_m_ (*Greene, Cao, Schanne, & Bartelt, 2002*; *Nelson et al., 2004*; *Song, Liu, et al., 2016*). The fragments were subcloned into the *Bam*HI site of the plasmid prg3-AMAI-NotI to generate pAMA1-P_gpdA_-Aeq and pAMA1-P_gpdA_-mt-Aeq, respectively.

For monitoring the calcium level in the vacuole, the plasmid pAMA1-P_gpdA_-CpyA-Aeq was generated as follows: PCR was performed using the primers ClaI-cpyA-R and ClaI-cpyA-F to generate a *cpyA’* ORF fragment. The fragment was subcloned into the *Cla*I site of pBARGPE, which has a P_gpdA_ promoter in front of its *Cla*I site, to generate the plasmid pBARGPE-P_gpdA_-cpyA. The P_gpdA_-cpyA fragment was amplified from the pBARGPE-P_gpdA_-cpyA with Ama1-BamHI-gpd-F and CpyA-linker-R primers and then fused with Aeq-TrpC fragment generated using Linker-Aeq-F and Ama1-BamHI-trpC-R to yield fusion fragment P_gpdA_-cpyA-Aeq. The pAMA1-P_gpdA_-CpyA-Aeq plasmid was generated by ligating P_gpdA_-cpyA-Aeq into prg3-AMAI-NotI.

pAMA1-P_gpdA_-CpyA-GFP/RFP was generated as follows: Fusion PCR using primers Ama1-BamHI-gpd-F and Ama1-BamHI-GFP/RFP-R, was performed to fuse the P_gpdA_-cpyA with GFP/RFP fragment amplified by primers Linker-GFP/RFP-F and Ama1-BamHI-GFP/RFP-R. Then, the fusion fragment was ligated into the *Bam*HI site of the plasmid prg3-AMAI-NotI, generating pAMA1-P_gpdA_-CpyA-GFP/RFP.

For indicating the vacuole, the plasmid pAMA1-P_gpdA_-CpyA-GFP/RFP was generated as follows: Fusion PCR using primers Ama1-BamHI-gpd-F and Ama1-BamHI-GFP/RFP-R, was performed to fuse the P_gpdA_-cpyA with GFP/RFP fragment amplified by primers Linker-GFP/RFP-F and Ama1-BamHI-GFP/RFP-R. Then, the fusion fragment was ligated into the *Bam*HI site of the plasmid prg3-AMAI-NotI to generate pAMA1-P_gpdA_-CpyA-GFP/RFP.

For visualization of cell nucleus, the pAMA1-P_gpdA_-RFP-H2A was generated as follows: The P_gpdA_-RFP-H2A fragment was amplified from the pBARGPE-P_gpdA_-RFP-H2A with primers Ama1-BamHI-gpd-F and Ama1-BamHI-trpC-R and then was subcloned into the *Bam*HI site of the plasmid prg3-AMAI-NotI to generate pAMA1-P_gpdA_-RFP-H2A.

For visualization of ER, the plasmid pAMA1-P_gpdA_-erg11A-GFP was generated as follows: using primers Ama1-BamHI-gpd-F and Erg11A-R, the P_gpdA_-erg11A fragment was amplified from the CX23 (*Song, Zhai, et al., 2016*). Using primers Erg11A-GFP-F and Ama1-BamHI-GFP-R, the GFP was amplified. Then, P_gpdA_-erg11A and GFP were fused together with primers Ama1-BamHI-gpd-F and Ama1-BamHI-GFP-R. Subsequently, the fusion fragment was subcloned into the *Bam*HI site of the plasmid prg3-AMAI-NotI to generate pAMA1-P_gpdA_-erg11A-GFP.

For visualization of mitochondria, the plasmid pAMA1-MrsA-RFP was generated as follows: Using primers Ama1-BamHI-MrsA-F and MrsA-R, the MrsA fragment containing *mrsA* promoter and ORF without stop codon was amplified from the *A. fumigatus* genomic DNA (gDNA). The RFP was amplified with primers MrsA-RFP-F and Ama1-BamHI-RFP-R. Then, MrsA fragment and RFP were fused together with primers Ama1-BamHI-MrsA-F and Ama1-BamHI-RFP-R. Subsequently, the fusion fragment was subcloned into the *Bam*HI site of the plasmid prg3-AMAI-NotI to generate pAMA1-MrsA-RFP.

For visualization of Golgi apparatus pAMA1-P_gpdA_-mRFP-PH^OSBP^ was generated as follows: The pleckstrin homology domain of the human oxysterol binding protein (PH^OSBP^) was PCR-amplified with primers gpdA-RFP-F and BamHI-PH^OSBP^-R and then fused with P_gpdA_ promoter (PCR-amplified with Ama1-BamHI-gpd-F and Gpd-R) using primers Ama1-BamHI-gpd-F and BamHI-PH^OSBP^-R. The fusion fragment was subcloned into *Bam*HI site of the plasmid prg3-AMAI-NotI to pAMA1-P_gpdA_-mRFP-PH^OSBP^.

For measuring the level of autophagy, plasmid pAMA1-P_gpdA_-GFP-Atg8 overexpressing GFP-Atg8 was generated as follows: PCR, using the joint primers Ama1-BamHI-gpd-F/Gpd-GFP-F/Atg8-F and Gpd-R/Atg8-GFP-R/Ama1-BamHI-Atg8-R, were performed to generate P_gpdA_ promoter/GFP/Atg8(ORF) fragments. Next, the fusion fragments P_gpdA_-GFP-Atg8 were generated by fusion PCR with primers Ama1-BamHI-gpd-F and Ama1-BamHI-Atg8-R. The fusion fragments were subcloned into the *Bam*HI site of the plasmid prg3-AMAI-NotI to generate pAMA1-P_gpdA_-GFP-Atg8. The plasmid pAMA1-P_atg8_-GFP-Atg8 expressing GFP-Atg8 under *atg8* native promoter was generated as follows: PCR, using the joint primers Ama1-Bam1-atg8(p)-F/Atg8(p)-GFP-F/ and Atg8-promoter-R/Ama1-BamHI-Atg8-R, was used to generate *atg8* promoter/GFP-Atg8 fragments. Next, the two fragments were fused together with primers with primers Ama1-Bam1-atg8(p)-F and Ama1-BamHI-Atg8-R. The fusion fragment was subcloned into the *Bam*HI site of prg3-AMAI-NotI to generate pAMA1-P_atg8_-GFP-Atg8.

For assessing the co-localization of CpyA-RFP and GFP-Atg8, the plasmid pAMA1-P_gpdA_-CpyA-RFP-P_gpdA_-GFP-Atg8 was generated as follows: The fragment P_gpdA_-CpyA-RFP/P_gpdA_-GFP-Atg8 was amplified with primers pairs Ama1-BamHI-gpd-F and TAA-RFP-R/RFP-gpdA-F and Ama1-BamHI-Atg8-R. Next, the two fragments were subcloned into prg3-AMAI-NotI with the One Step Cloning Kit (Vazyme, C113) forming pAMA1-P_gpdA_-CpyA-RFP-P_gpdA_-GFP-Atg8.

For overexpressing *pmcA/pmcB*, plasmid pBARGPE-hph-P_gpdA_-pmcA/pmcB overexpressing *pmcA/pmcB* were generated as follows: PCR, using the joint primers ClaI-pmcA/pmcB-F and ClaI-pmcA/pmcB-R, were used to generate pmcA/pmcB fragment that include the complete ORF and 3’UTR of the indicated genes. Next, the three fragments were, respectively, subcloned into the *Cla*I site of the plasmid to generate overexpression plasmids pBARGPE-hph-P_gpdA_-pmcA/pmcB, which contain a selective marker gene *hph* to resist hygromycin.

For amplifying the repair template gfp-hph, the plasmid p-zero-gfp-hph was generated as follows: the selectable marker *hph* was amplified by PCR using the primers HPH-F and HPH-R, and then cloned into the pEASY-Blunt vector, generating a plasmid p-zero-hph. Primers NOTI-gfp-F and NOTI-gfp-R were used to generate a GFP fragment. This fragment was then cloned into the *Not*I site of p-zero-hph to generate p-zero-gfp-hph.

For amplifying the repair template ptrA-P_gpdA_/ptrA-P_gpdA_-gfp, the plasmid p-zero-ptrA-P_gpdA_-gfp was generated as follows: the selectable marker *ptrA* was amplified by PCR using the primers ptrA-F and ptrA-R, and then cloned into the pEASY-Blunt vector, generating a plasmid p-zero-ptrA. Next, the plasmid was linearized with primer zero-F and zero-R. PCR, using primers zero-PgpdA-F/ PgpdA-GFP-F and PgpdA-R/zero-GFP-R, were used to amplify P_gpdA_ promoter/gfp fragments. The two fragments were fused with primers zero-PgpdA-F and zero-GFP-R, generating P_gpdA_-gfp fragment. Finally, the fusion fragment was subcloned into the linearizing p-zero-ptrA, yielding the plasmid p-zero-ptrA-P_gpdA_-gfp.

For amplifying the repair template hph-P_gpdA_, the plasmid p-hph-PgpdA was generated as follows: PCR, using primer NotI-PgpdA-F and NotI-TtrpC-R, was used to amplify *gpdA* promoter (P_gpdA_). Then, the *gpdA* promoter was ligated into the *Not*I site of p-zero-HPH, generating p-hph-PgpdA.

For expressing the N-terminus (300 aa) of CchA, the plasmid pAMA1-P_gpdA_-GFP-CchA^N300^ was generated as follows: The P_gpdA_-GFP fragment was obtained by PCR with primers, Ama1-BamHI-gpd-F and CchA-GFP-R from p-zero-ptrA-P_gpdA_-gfp; The CchA^N300^ fragment containing 900 bp of *cchA* ORF 5’ was amplified by PCR with primer CchA-F and Ama1-BamHI-CchA^N300^-R. Then, the two fragments were subcloned into prg3-AMAI-NotI with the One Step Cloning Kit (Vazyme, C113), generating the plasmid pAMA1-P_gpdA_-GFP-CchA^N300^.

For live animal bioluminescence imaging, the plasmid pBARGPE-pyr4-P_gpdA_-Luc expressing Luciferase was generated as follows: PCR, using the joint primers ClaI-Luc-F and ClaI-Luc-R, were used to generate *luc* (Luciferase gene) ORF. Next, the Luc fragments was subcloned into the *Cla*I site of the plasmid pBARGPE-pyr4, yielding pBARGPE-pyr4-P_gpdA_-Luc.

All the primers and primer annotations were given in supplementary data Table S2.

### Gene-editing with MMEJ-CRISPR in *A. fumigatus,* transformation and strain verification

For editing the *A. fumigatus* gene, the MMEJ-CRISPR system was used as described in our previous published papers (*Zhang & Lu, 2017*; *Zhang, Meng, Wei, & Lu, 2016*). sgRNA targeted to the appropriate site of the target gene was synthesized *in vitro* using the MEGAscript T7 Kit (Life Technologies, AM1333). The corresponding repair template (DNA) with microhomology arms was amplified by PCR. Then, the repair template fragments and sgRNA were cotransformed into a Cas9-expressing *A. fumigatus* recipient strain. The primers and annotations for sgRNAs and repair templates are listed in Table S2. Transformation procedures were performed as previously described. Transformants were selected in medium lacking uridine or uracil or in the presence of 150 μg/ml hygromycin B (Sangon) or 0.1 μg/ml pyrithiamine (Sigma). For the recycling usage of the selectable marker *pyr4*, 1 mg/mL 5-FOA was used for screening recipient strains. All primers used are listed in the supplementary data Table S2. All transformant isolates were verified by diagnostic PCR analysis using mycelia as the source of DNA. Primers were designed to probe upstream and downstream of the expected cleavage sites as labeled in Figure S1D.

### Microscopy observation

For microscopic observation of hyphae morphology, fresh conidia were inoculated onto sterile glass coverslips overlaid with 1 mL of liquid glucose media with or without 10 mM CaCl_2_. Strains were cultivated on the coverslips at 37°C for 14 h before observation. The coverslips with hyphae were gently washed with PBS three times. Differential interference contrast (DIC) and green/red fluorescent images of the cells were collected with a Zeiss Axio Imager A1 microscope (Zeiss, Jena, Germany). To observe the fluorescence localization of GFP/RFP-tagged proteins, strains were grown in liquid medium at 37°C with shaking at 200 rpm for indicated time, and then the mycelium pellet sandwiched between the microscope slide and coverslips was observed directly by fluorescence microscopy.

### Measurement of the free Ca^2+^ concentration ([Ca^2+^])

Measurement of the free Ca^2+^ concentration was performed as described previously with some modifications *(Nelson et al., 2004;* Y. *Zhang et al., 2016*). The strains expressing Aeq/Mt-Aeq/CpyA-Aeq were cultured for 2 days to form fresh spores. The spores were filtered through nylon cloth and washed 10 times in distilled deionized water. One million (10^6^) spores in 100 μL liquid MM were inoculated into each well of a 96-well microtiter plate (Thermo Fischer). The plate was incubated at 37°C for 24 h. The medium was then removed gently with a pipette, and the cells in each well were washed twice with 150 µL PGM (20 mM PIPES pH 6.7, 50 mM glucose, 1 mM MgCl_2_). Aequorin was reconstituted by incubating mycelia in 100 μL PGM containing native coelenterazine (2.5 μM) (Sigma-Aldrich, C-7001) at 4°C for 4 h in the dark. Then, mycelia were washed twice with 150 µL PGM and allowed to recover to room temperature for 1 h. Luminescence was measured with an LB 960 Microplate Luminometer (Berthold Technologies, Germany), which was controlled by a dedicated computer running the MikroWin 2000 software. At the 20-s time point of luminescence reading, 10 mM CaCl_2_ was applied as a stimulant. At the end of each experiment, the active aequorin was completely discharged by permeabilizing the cells with 20% (vol/vol) ethanol in the presence of an excess of calcium (3 M CaCl_2_) to determine the total aequorin luminescence of each culture. The conversion of luminescence (relative light units [RLU]) into [Ca^2+^] was performed with using Excel 2019 software (Microsoft). Input data were converted using the following empirically derived calibration formula: pCa = 0.332588 (-log k) + 5.5593, where k is luminescence (in RLU) s^-1^/total luminescence (in RLU).

### *In vitro* phosphorylation analysis

GFP-CchA^N300^-expressing *A. fumigatus* were grown in liquid MM for 36 with shaking at 200 rpm. Then, the mild buffer (10 mM Tris-HCl, pH 7.5, 150 mM NaCl, 0.5 mM EDTA, 0.01% Triton X-100, 1 mM DTT, 1 mM PMSF, and 1:100 protease inhibitor cocktail) was used to extract the total proteins. The GFP-CchA were enriched with a GFP-Trap kit (ChromoTek). After enrichment, samples were separated on 10% SDS-PAGE. The gel bands corresponding to the targeted protein GFP-CchA were excised from the gel. The gel pieces were destained 3 times with 125 mM NH_4_HCO_3_/50% acetonitrile. Then, proteins were reduced with 10 mM DTT/125 mM NH_4_HCO_3_ at 56°C for 30 min and alkylated with 55 mM iodoacetamide/125 mM NH_4_HCO_3_ for 30 min in the dark. After washing twice with 50% acetonitrile, the gel pieces were dried in a vacuum centrifuge. Then, 200 ng trypsin in 25 mM NH_4_HCO_3_ was added, and the digestion was maintained at 37°C overnight. After digestion, 0.5% formic acid/50% acetonitrile was added to extract peptides from the gel pieces. The extraction procedure was repeated, and the three extracts were combined and dried by vacuum centrifugation. The peptide samples were dissolved in 2% acetonitrile/0.1% formic acid and analyzed using a TripleTOF 5600+ mass spectrometer coupled with the Eksigent nanoLC System (SCIEX, USA). Peptide was loaded onto a C18 trap column (5 μm, 100 μm×20 mm) and eluted at 300 nL/min onto a C18 analytical column (3 μm, 75 µm×150 mm) over a 60-min gradient. The two mobile phases were buffer A (2% acetonitrile/0.1% formic acid/98% H_2_O) and buffer B (98% acetonitrile/0.1% formic acid/2% H_2_O). For information dependent acquisition (IDA), survey scans were acquired in 250 ms, and 40 product ion scans were collected in 50 ms/per scan. MS1 spectra were collected in the range 350-1500 m/z, and MS2 spectra were collected in the range of 100-1500 m/z. Precursor ions were excluded from reselection for 15 s. Phosphorylation site identification was achieved by searching the obtained MS spectra files using Mascot Daemon software.

### Proteomics

The Proteomics experiment was performed at Hangzhou PTM Biolabs Co., Ltd. as a commercial service. In brief, lysis buffer (8 M urea, 1% Triton-100, 10 mM dithiothreitol, and 1% Protease Inhibitor Cocktail (Calbiochem) was used to extract the total protein. Proteins were digested with trypsin (Promega) to be peptides and then processed according to the manufacturer’s protocol for TMT kit. The labeled tryptic peptides were fractionated by HPLC using a Thermo Betasil C18 column (5 μM particles, 10 mm ID, 250 mm length). After fractioning, the tryptic peptides were analyzed with LC-MS/MS system. The MS/MS data processing were performed using Maxquant search engine (v.1.5.2.8). Tandem mass spectra were searched against *Aspergillus* UniProt database concatenated with reverse decoy database. Proteins with change ratios significantly different from general protein variation (1.50 < Δ*cnaA*/wild-type < 0.50) were analyzed for GO terms and KEGG pathway analysis using the UniProt-GOA database (http://www.ebi.ac.uk/GOA/) and the KEGG database, respectively.

### Virulence Assay

For *G. mellonella* infection model, *G. mellonella* larva, in groups of 10, were injected through one of the hind pro-legs with 10 μl of PBS containing 8 x 10^7^ conidia of the respective strain. Untreated larvae and larvae injected with 10 μl of PBS served as controls. Larvae were incubated at 37°C in the dark and monitored daily for up to 10 days. The significance of survival data was evaluated using Kaplan-Meier survival curves and analyzed with the log-rank (Mantel Cox) test on GraphPad Prism software. Differences were considered significant at *p* values < 0.05.

For mouse infection model challenged with *A. fumigatus* conidia, ICR mice (6–8 weeks old, male, 25 ± 5 g) were purchased from Beijing Vital River Laboratory Animal Technology Co., Ltd. The mice were randomized into four groups (n = 10 each). Mice were immunosuppressed on day -3 and -1 with cyclophosphamide (150 mg/kg) and on day -1 with hydrocortisone acetate (40 mg/kg). On day 0, mice were anesthetized with 1% pentobarbital sodium (10 ml/kg) and infected intratracheally with a 50-μL slurry containing 5 × 10^6^ conidia or 50 μL of PBS as the control according to previously described methods (REF Li et al. (2014) and Zhang et al. (2015)). Cyclophosphamide (150 mg/kg) was injected every 3 days after infection to maintain immunosuppression. Survival was monitored two weeks after inoculation. Lung tissue was processed for histology, and Grocott’s methenamine silver stain and periodic acid-Schiff (PAS) staining were performed. For mouse infection model challenged with *A. fumigatus* hyphae, the mycelia of each strain were homogenized into hyphae fragments using tissue grinding and filtered with lens-cleaning paper in sterile normal saline. The optical density at 600 nm of the 10-fold dilutions of each group were finally adjusted to 0.5. The ICR mice described as above (4 groups, n=10 each) were immunosuppressed on day -3 and -1 with cyclophosphamide (150 mg/kg). On day 0, mice were anesthetized with pentobarbital sodium and then infected intratracheally with a 50-µl hyphal suspension. Cyclophosphamide (150 mg/kg) was injected every 3 days to maintain immunosuppression. Survival was monitored 10 days after inoculation. To directly monitor the infection status in the hyphal infection model, luciferase-expressing *A. fumigatus* strains were used to infect mice, and live mouse bioluminescence imaging was performed. The mice (4 groups, n=5 each) were immunosuppressed and infected as in the above description. The infected mice were introduced intraperitoneally with 100 µl D-luciferin in PBS (33.3 mg/ml) 10 min before bioluminescence imaging. Mice were anesthetized using 2.5% isoflurane with a constant flow and then placed in an IVIS Lumina XR system chamber. Scanning was performed on day 4. The region was cropped in the chest area from each animal, and the total bioluminescence signal intensity of the lung region was extracted and analyzed using Living Image 4.2 (Caliper Life Sciences).

### Statistics

Data were given as means ± SD. The SD was from at least three biological replicates. Statistical significance was estimated with Graphpad Prism 7 using Student’s *t*-test or log-rank test. P-values less than 0.05 were considered statistically significant.

## Acknowledgments

This work was financially supported by the National Key Research and Development Program of China (grant 2019YFA0904900), the National Natural Science Foundation of China (grants NSFC31861133014 and 31770086), the Program for Jiangsu Excellent Scientific and Technological Innovation team (grant 17CXTD00014), the Priority Academic Program Development of Jiangsu Higher Education Institutions, and National Natural Science Foundation of China (31900404) and Natural Science Foundation of the Jiangsu Higher Education Institutions of China (19KJB180017) to Y.Z. We thank Gustavo H. Goldman (University of São Paulo) for kindly providing helpful comments on the manuscript.

## Author contributions

C.Z., Y.Z. and L.L., conceived the study; C.Z., Y.R., H.G., and L.G. performed the experiments; and C.Z., Y.Z. and L.L. analyzed and interpreted the data and wrote the manuscript with input from all authors.

## Supplement Figures

**Figure S1.**
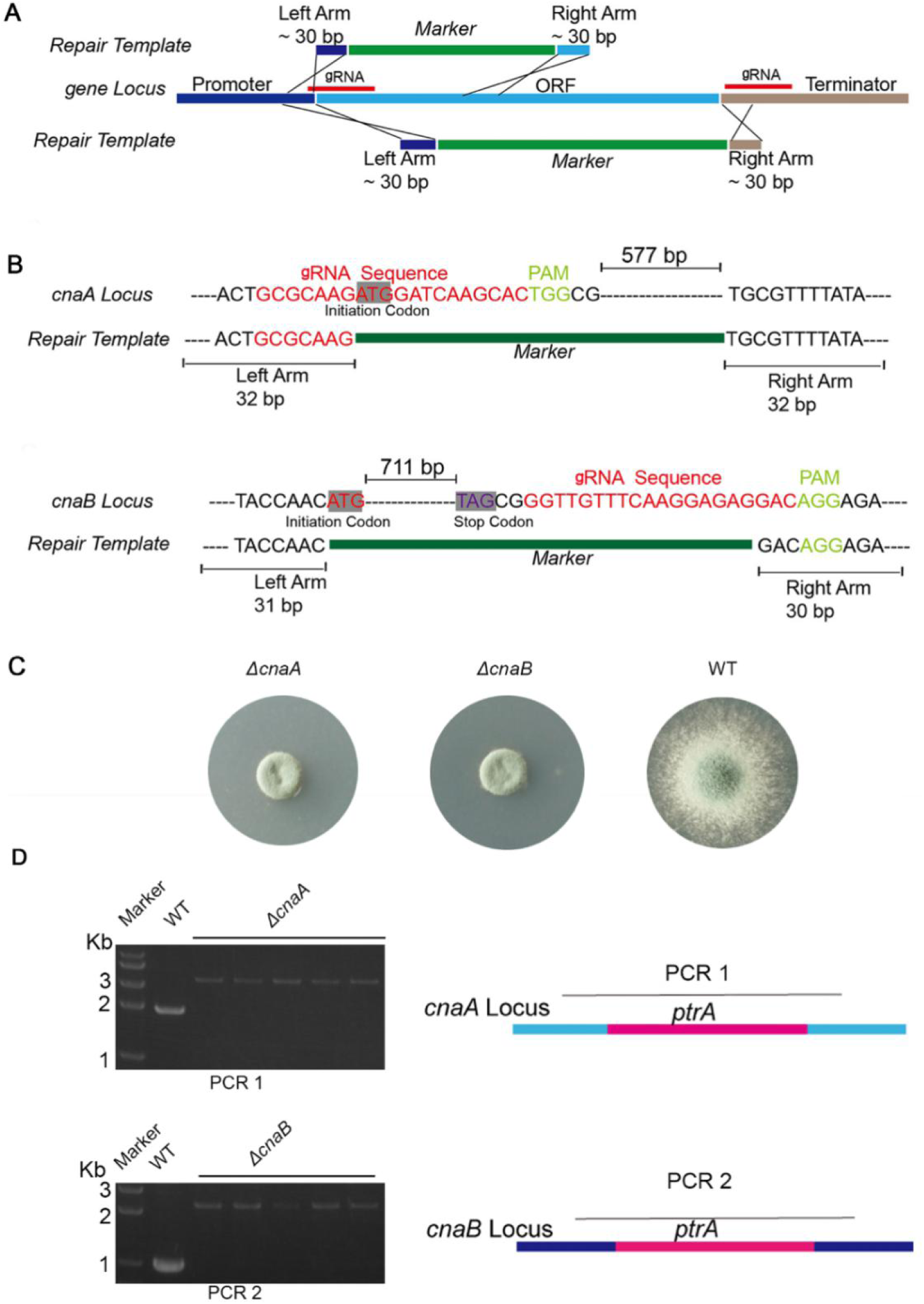
Constructions of *cnaA* and *cnaB* deletion mutants in *A. fumigatus* via CRISPR-Cas9 system. A. Schematic illustration of gRNA location and repair template for deleting genes via the CRISPR-Cas9 system. B. Details of gRNA and the repair template for *cnaA* and *cnaB* deletion. C. The colony morphologies of Δ*cnaA,* Δ*cnaB*, and wild-type *A. fumigatus*. D. Diagnostic PCR for *ptrA* integration in the corresponding gene’s ORF. Each pair of diagnostic primers was complementary to upstream and downstream sequences flanking the deletion fragment.

**Figure S2.**
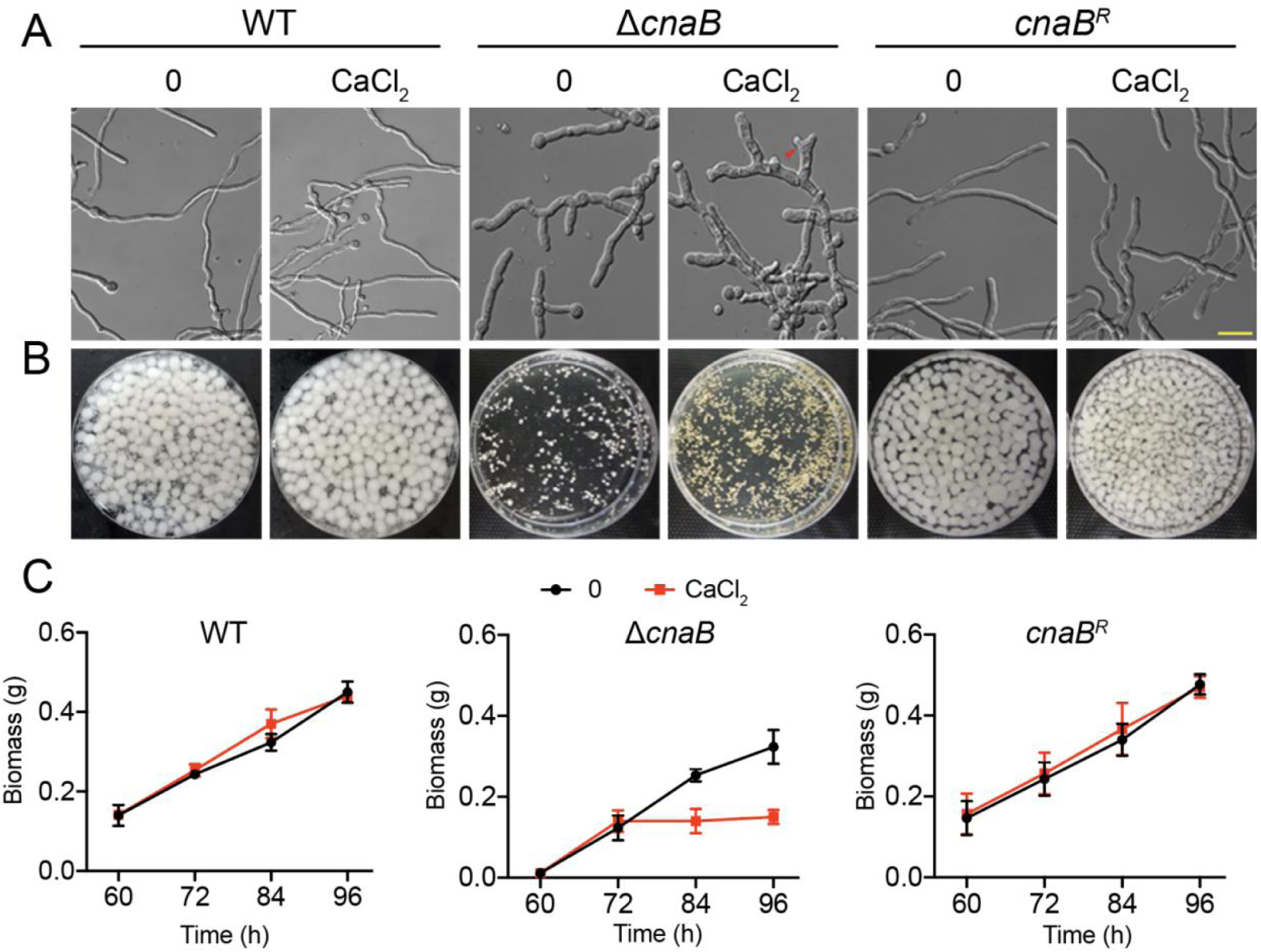
Phenotypic characterization of the Δ*cnaB* mutant. A. Differential interference contrast (DIC) images of hyphae grown in liquid MM with or without 10 mM CaCl_2_ at 37°C in stationary culture for 14 h. Scale bar represents 10 μm. B. The morphology of mycelial pellets of the indicated strains grown in MM with or without 10 mM CaCl_2_ at 37°C for 60 h. C. Quantification of biomass production for the wild-type, Δ*cnaB*, and *cnaB^R^* complementary strains grown in MM with or without 10 mM CaCl_2_ at different time points (60, 72, 84, and 96 h).

**Figure S3.**
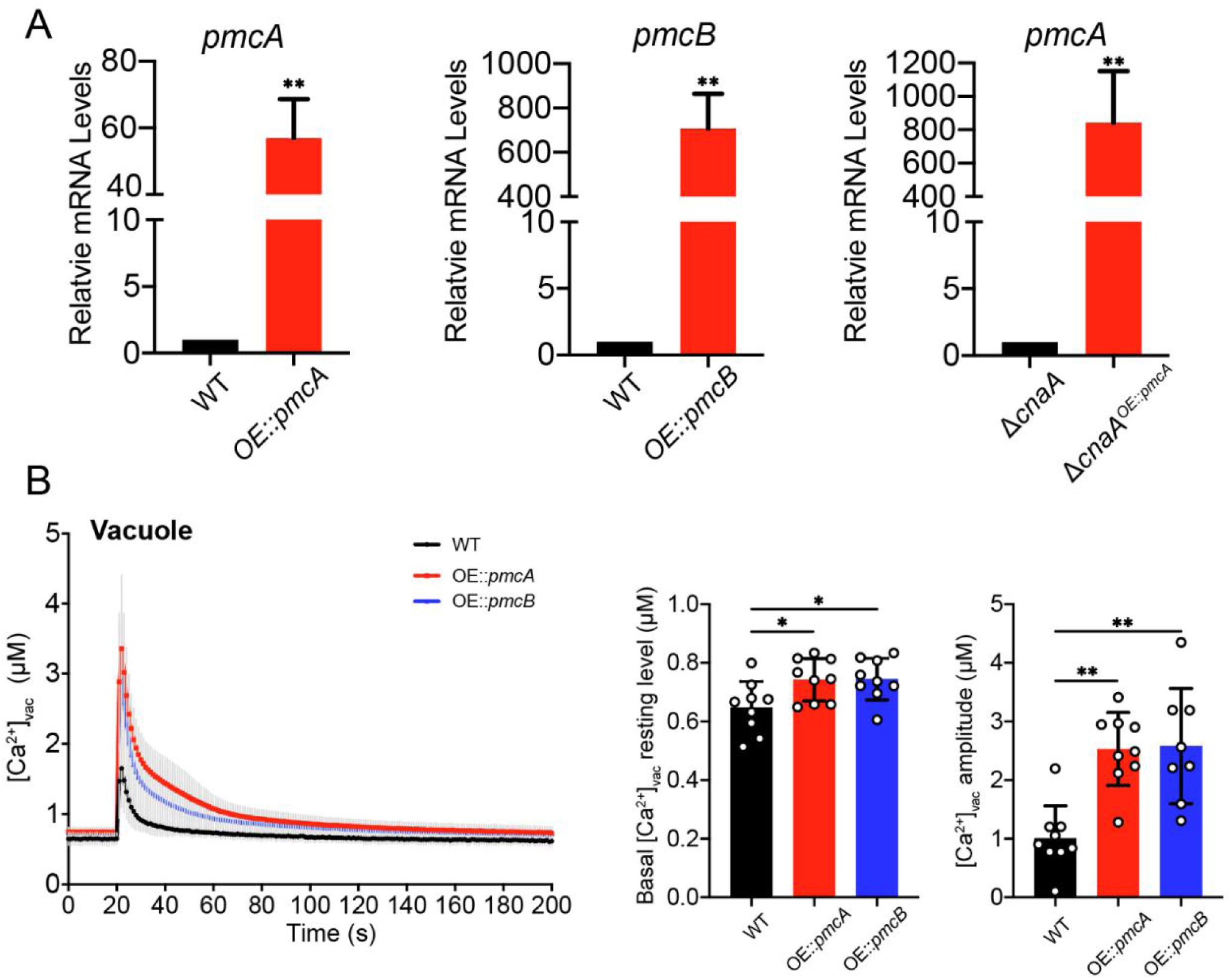
mRNA expression analysis of *pmcA* and *pmcB* in the wild-type and Δ*cnaA* background strains and calcium-induced [Ca^2+^]_vac_ transients in the wild-type, OE::*pmcA*, and OE::*pmcB* strains. A. Transcript levels of *pmcA/B* in WT and OE::*pmcA/*B (*pmcA/B*-overexpressed strains) and in Δ*cnaA* and Δ*cnaA*^OE::*pmcA*^ were determined by qRT-PCR. The indicated strains of *A. fumigatus* were incubated in MM for 36 h at 37°C. \*\**p*< 0.01. B. Comparison of resting and dynamic [Ca^2+^]_vac_ levels in the wild-type and *OE::pmcA/B* strains. Statistical significance was determined by Student’s *t*-test. \**p* < 0.05; \*\**p*< 0.01.

**Figure S4.**
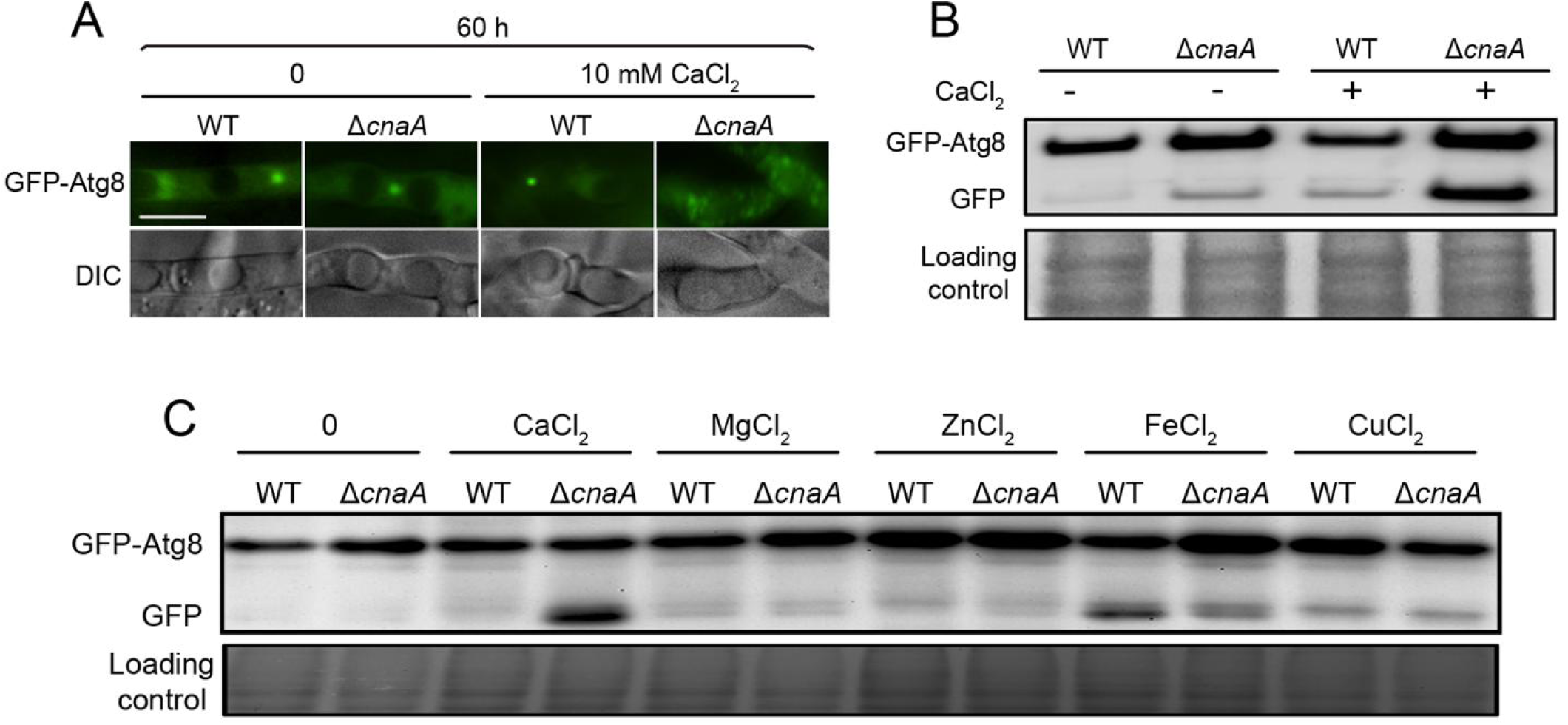
Fluorescence microscope observation and Western blots were used to examine the cleavage and localization of GFP-Atg8 under the control of the native *atg8* promoter (A and B) or the *gpdA* promoter (C). WT and Δ*cnaA* were cultured in MM with or without various divalent ions (10 mM CaCl_2_, 10 mM MgCl_2_, 10 mM ZnCl_2_, 2 mM FeCl_2_, or 0.2 mM CuCl_2_) for 60 h. Scale bar represents 5 μm.

**Figure S5.**
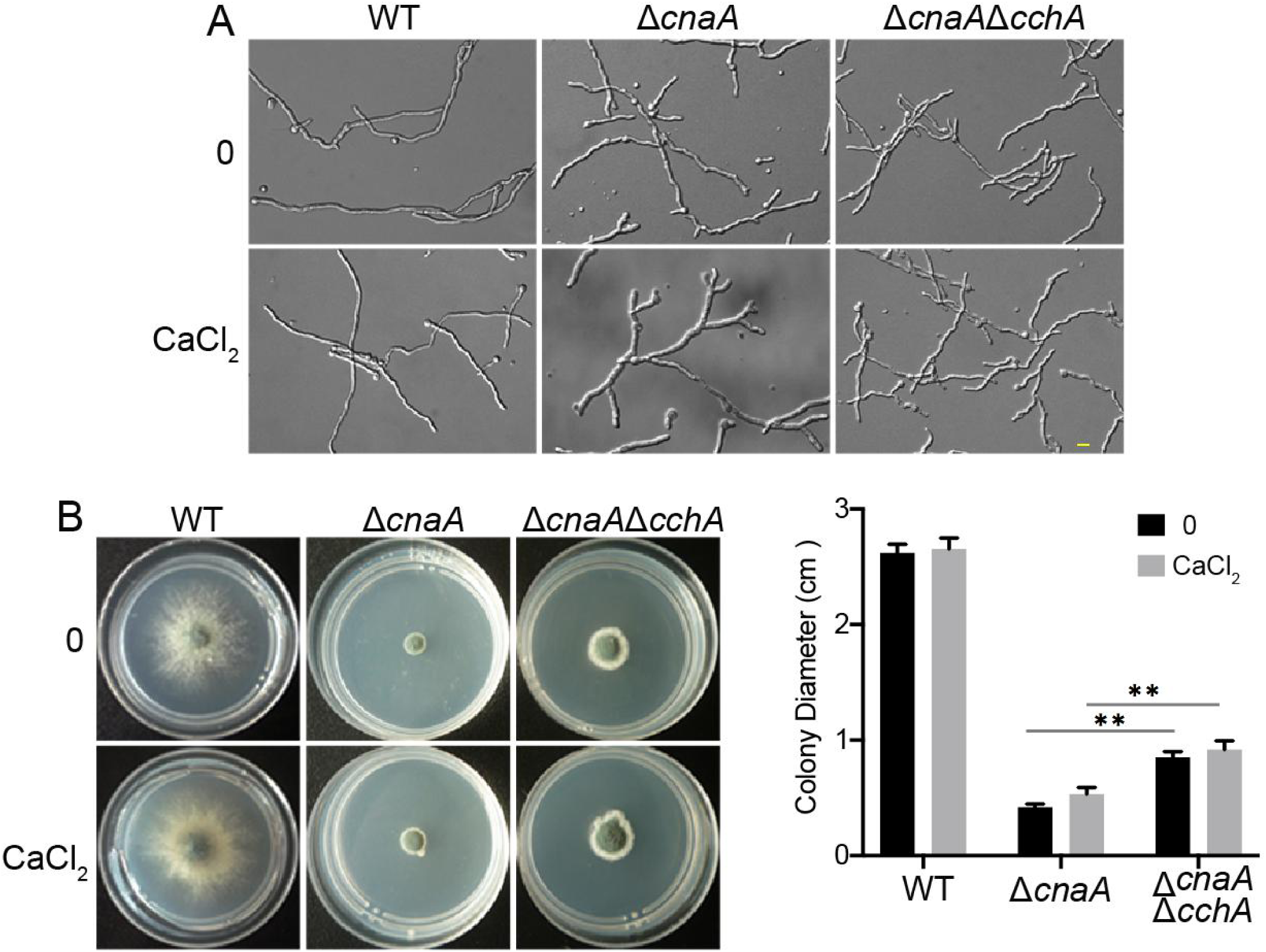
Growth morphologies of the Δ*cnaA*Δ*cchA* mutant. A. DIC images of hyphae grown in liquid MM with or without 10 mM CaCl_2_ at 37°C for stationary culture 14 h. Scale bar represents 10 μm. B. Colony morphologies of WT, Δ*cnaA* and Δ*cnaA*Δ*cchA* on solid MM or MM plus 10 mM CaCl_2_. The 1×10^4^ conidia were inoculated onto solid media for 48 h at 37°C. Statistical significance was determined by Student’s *t*-test. Statistical significance was determined by Student’s *t*-test. \*\**p* < 0.01.

**Figure S6.**
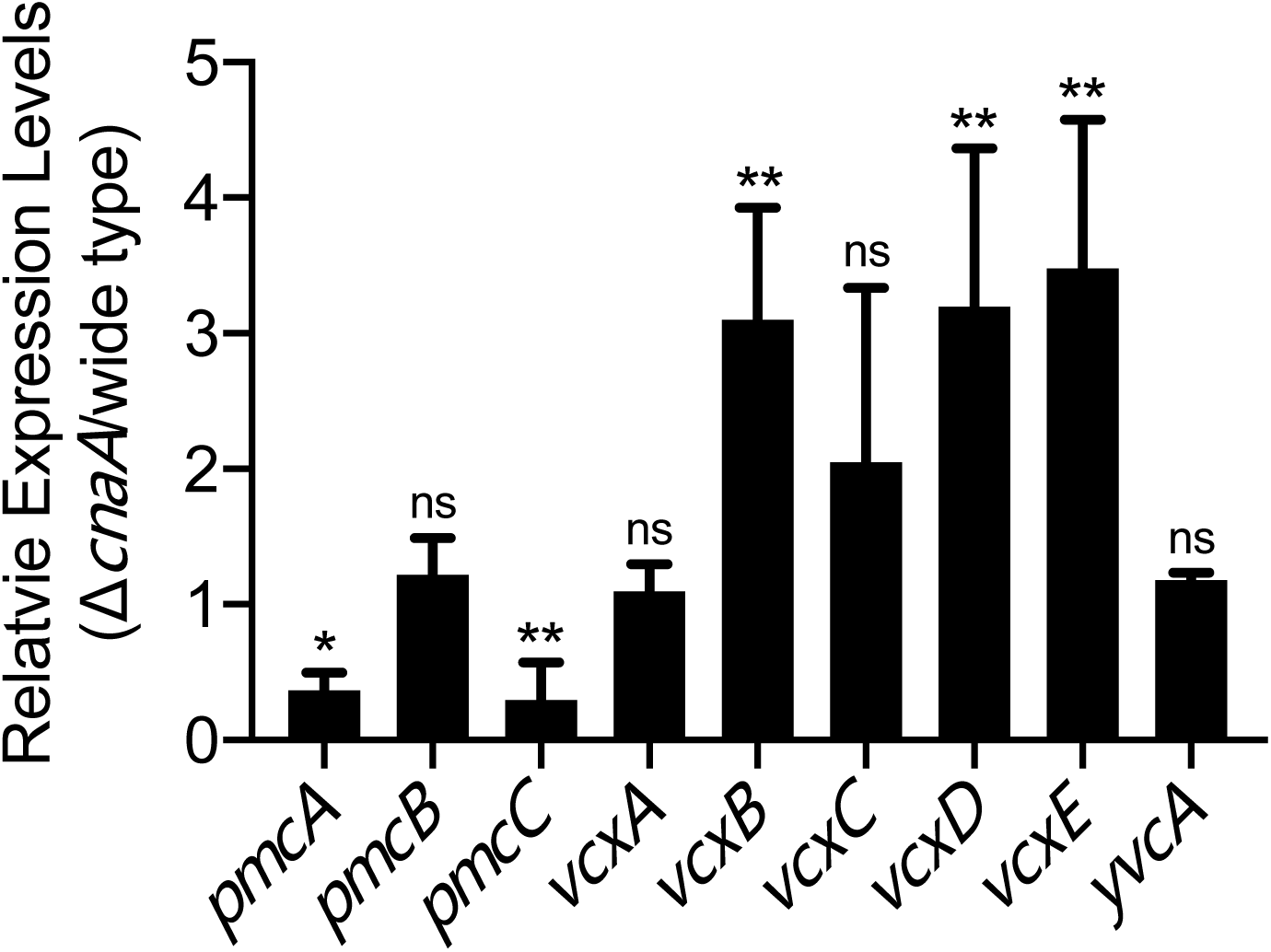
The transcript levels of *pmcA/B/C*, *vcxA/B/C/D/E*, and *yvcA* in the wild-type and Δ*cnaA* strains grown in MM for 36 h. Statistical significance was determined by Student’s *t*-test. \**p* < 0.05; \*\**p* < 0.01; ns, not significant.

**Figure S7.**
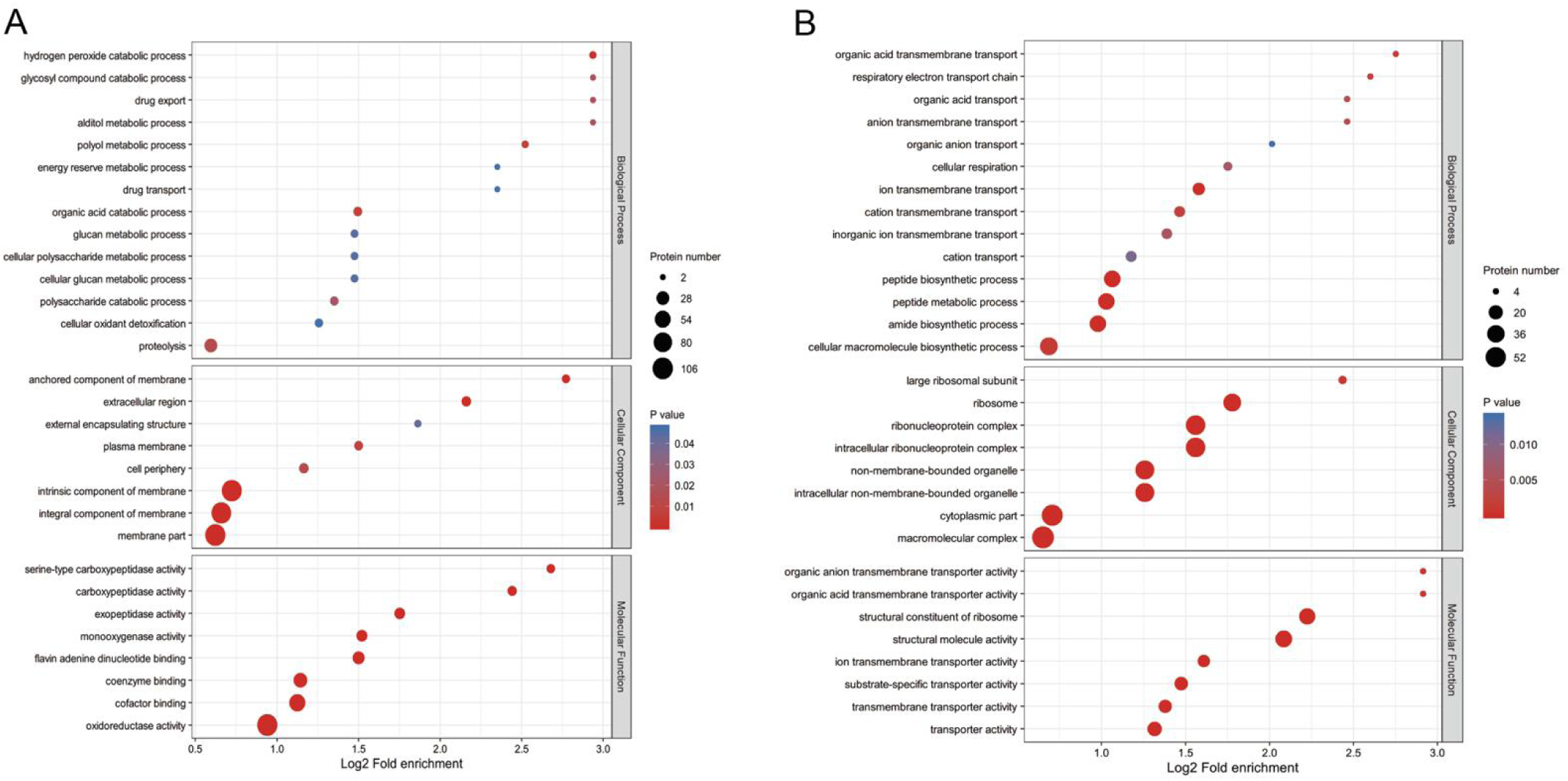
The enriched GO terms in the upregulated (A) and downregulated (B) proteins with greater than 1.5-fold changes in the Δ*cnaA* mutant compared to the wild-type. GO annotation was derived from the UniProt-GOA database (http://www.ebi.ac.uk/GOA/).

**Figure S8.**
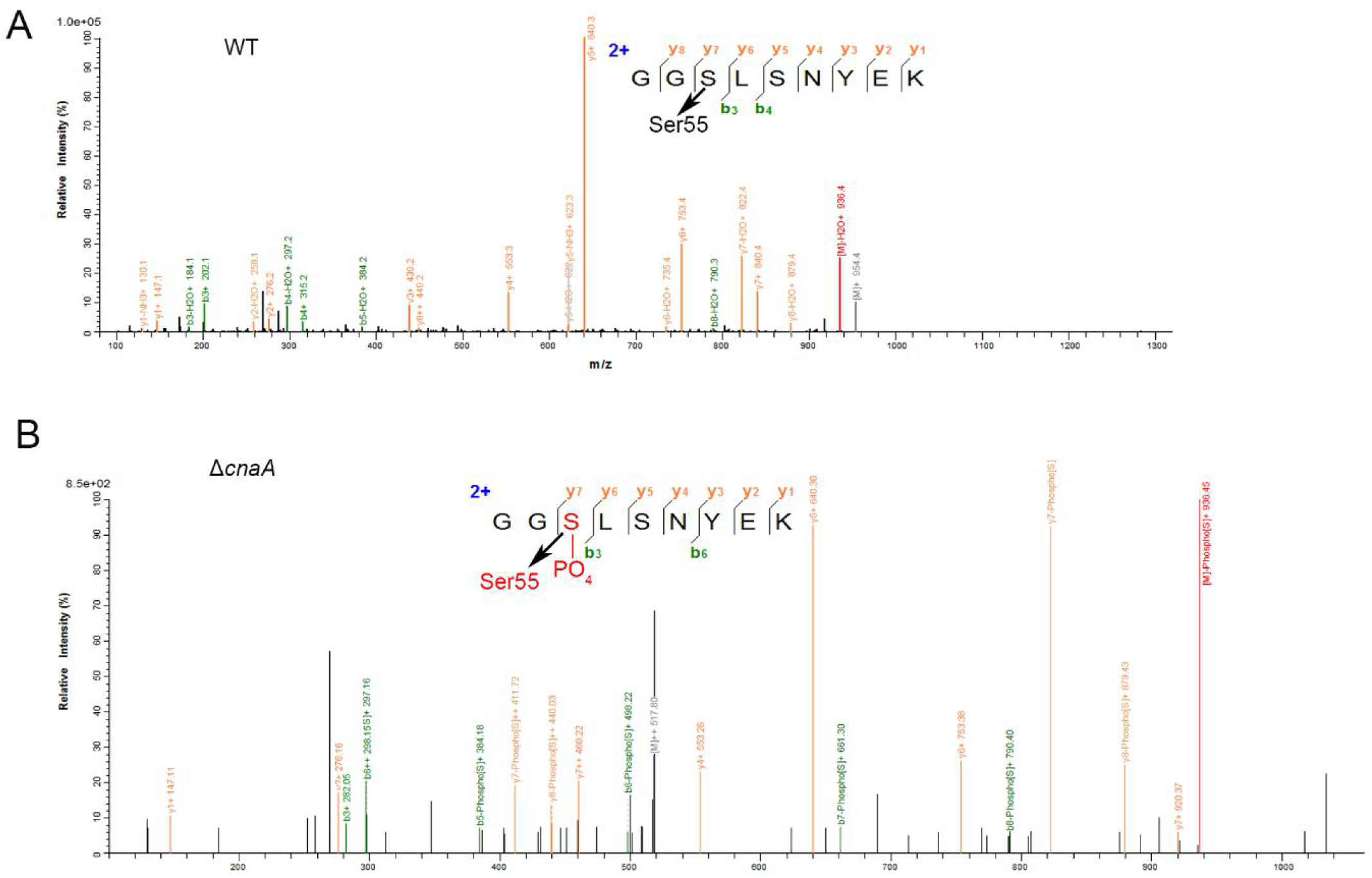
CchA is dephosphorylated at S55 by calcineurin *in vivo*. A and B. Tandem mass spectra of GGSLSNYEK from CchA reveal one unique phosphorylated serine residue, S55, in Δ*cnaA.* The presence of identified C-terminal (y) and N-terminal (b) product ions are indicated within the peptide sequence. The b and y ions are indicated with red and green colors, respectively.

