## Supplemental Table S1 for "When and why an essential popular mineral—calcium—is an effective antifungal adjuvant against the human pathogen?"

Table S1. A list of *A. fumigatus* strains used in this study.

| Strains | Genotype And Source |
| --- | --- |
| ZC03/WT | Δ*ku80; pyrG1;* *AMA1::P_gpdA_::Cas9::pyr4;* from A1160 transformed by FM-6 |
| Δ*cnaA* | ZC03；Δ*cnaA::ptrA* |
| Δ*cnaB* | ZC03；Δ*cnaB::ptrA* |
| *cnaA^R^* | Δ*ku80; pyrG1;* *AMA1::P_gpdA_::Cas9::pyr4;* Δ*cnaA::ptrA; cnaA::hph* |
| *cnaB^R^* | Δ*ku80; pyrG1;* *AMA1::P_gpdA_::Cas9::pyr4;* Δ*cnaB::ptrA; cnaB::hph* |
| Δ*cnaA*( uridine and uracil auxotroph ) | Δ*ku80; pyrG1;* Δ*cnaA::ptrA;* It was generated by deleting FM-6 by 5-FOA under Δ*cnaA* background*.* |
| WT^cyt^ | Δ*ku80; pyrG1;* *AMA1::P_gpdA_::Aeq::pyr4* |
| Δ*cnaA^cyt^* | Δ*ku80; pyrG1;* Δ*cnaA::ptrA;* *AMA1::P_gpdA_::Aeq::pyr4* |
| *cnaA^Rcyt^* | Δ*ku80; pyrG1;* Δ*cnaA::ptrA; cnaA::hph; AMA1::P_gpdA_::Aeq::pyr4* |
| WT^mt^ | Δ*ku80; pyrG1;* *AMA1::P_gpdA_::mt-Aeq::pyr4* |
| Δ*cnaA^mt^* | Δ*ku80; pyrG1;* Δ*cnaA::ptrA;AMA1::P_gpdA_::mt-Aeq::pyr4* |
| *cnaA^Rmt^* | Δ*ku80; pyrG1;* Δ*cnaA::ptrA; cnaA::hph; AMA1::P_gpdA_::mt-Aeq::pyr4* |
| WT^cpyA-gfp^ | Δ*ku80; pyrG1; AMA1::P_gpdA_::cpyA-GFP::pyr4* |
| *gfp-pmcA*  *cpyA-rfp* | Δ*ku80; pyrG1; P_gpdA_::gfp-pmcA::ptrA; AMA1::P_gpdA_::cpyA-RFP::pyr4* |
| WT^vac^ | Δ*ku80; pyrG1;* *AMA1::P_gpdA_::cpyA-Aeq::pyr4* |
| *OE::pmcA^vac^* | Δ*ku80; pyrG1;* *AMA1::P_gpdA_::cpyA-Aeq::pyr4; P_gpdA_::pmcA::hph* |
| *OE::pmcB^vac^* | Δ*ku80; pyrG1;* *AMA1::P_gpdA_::cpyA-Aeq::pyr4; P_gpdA_::pmcB::hph* |
| Δ*cnaA^vac^* | Δ*ku80; pyrG1;* Δ*cnaA::ptrA; AMA1::P_gpdA_::cpyA-Aeq::pyr4* |
| *cnaA^Rvac^* | Δ*ku80; pyrG1;* Δ*cnaA::ptrA; cnaA::hph; AMA1::P_gpdA_::cpyA-Aeq::pyr4* |
| WT^rfp-H2A^ | Δ*ku80; pyrG1;* *AMA1::P_gpdA_::RFP-H2A::pyr4* |
| Δ*cnaA*^rfp-H2A^ | Δ*ku80; pyrG1;* Δ*cnaA::ptrA; AMA1::P_gpdA_::RFP-H2A::pyr4* |
| WT^erg11A-gfp^ | Δ*ku80; pyrG1;* *AMA1::P_gpdA_::erg11A-GFP::pyr4* |
| Δ*cnaA*^erg11A-gfp^ | Δ*ku80; pyrG1;* Δ*cnaA::ptrA; AMA1::P_gpdA_::erg11A-GFP::pyr4* |
| WT^rfp-ph-osbp^ | Δ*ku80; pyrG1;* *AMA1::P_gpdA_::RFP-PH^OSBP^::pyr4* |
| *ΔcnaA*^rfp-ph-osbp^ | Δ*ku80; pyrG1;* Δ*cnaA::ptrA;AMA1::P_gpdA_::RFP-PH^OSBP^::pyr4* |
| WT^mrsA-rfp^ | Δ*ku80; pyrG1;* *AMA1::P_gpdA_::MrsA-RFP::pyr4* |
| Δ*cnaA^mrsA-rfp^* | Δ*ku80; pyrG1;* Δ*cnaA::ptrA; AMA1::P_gpdA_::MrsA-RFP::pyr4* |
| WT^PgpdA-gfp-atg8^ | Δ*ku80; pyrG1;* *AMA1::P_gpdA_::GFP-Atg8::pyr4* |
| Δ*cnaA^PgpdA-gfp-atg8^* | Δ*ku80; pyrG1;* Δ*cnaA::ptrA; AMA1::P_gpdA_::GFP-Atg8::pyr4* |
| WT^PgpdA-gfp-atg8,cpyA-rfp^ | Δ*ku80; pyrG1;* *AMA1::P_gpdA_::CpyA-rfp::P_gpdA_::GFP-Atg8::pyr4* |
| Δ*cnaA^PgpdA-gfp-atg8,cpyA-rfp^* | Δ*ku80; pyrG1;* Δ*cnaA::ptrA; AMA1::P_gpdA_::CpyA-rfp::P_gpdA_::GFP-Atg8::pyr4* |
| WT^Patg8-gfp-atg8^ | Δ*ku80; pyrG1;* *AMA1::P_atg8_::GFP-Atg8::pyr4* |
| Δ*cnaA^Patg8-gfp-atg8^* | Δ*ku80; pyrG1;* Δ*cnaA::ptrA; AMA1::P_atg8_::GFP-Atg8::pyr4* |
| Δ*cnaA*Δ*atg2* | ZC03*;* Δ*cnaA::ptrA;* Δ*atg2::hph* |
| Δ*cnaA*Δ*atg2*^PgpdA-gfp-atg8,^*^cpyA-rfp^* | Δ*ku80; pyrG1;* Δ*cnaA::ptrA;* Δ*atg2::hph; AMA1::P_gpdA_::CpyA-rfp::P_gpdA_::GFP-Atg8::pyr4* |
| Δ*cnaA*Δ*atg2^PgpdA-gfp-atg8^* | Δ*ku80; pyrG1;* Δ*cnaA::ptrA;* Δ*atg2::hph; AMA1::P_gpdA_::GFP-Atg8::pyr4* |
| Δ*cnaA*Δ*cchA* | ZC03; Δ*cnaA::ptrA;* Δ*cchA::hph* |
| Δ*cnaA*Δ*cchA*^PgpdA-gfp-atg8,^*^cpyA-rfp^* | Δ*ku80; pyrG1;*Δ*cnaA::ptrA; ΔcchA::hph; AMA1::P_gpdA_::CpyA-rfp::P_gpdA_::GFP-Atg8::pyr4* |
| Δ*cnaA*Δ*cchA^PgpdA-gfp-atg8^* | Δ*ku80; pyrG1;* Δ*cnaA::ptrA;* Δ*cchA::hph; AMA1::P_gpdA_::GFP-Atg8::pyr4* |
| Δ*cnaA*Δ*cchA*^rfp-H2A^ | Δ*ku80; pyrG1;* Δ*cnaA::ptrA;* Δ*cchA::hph; AMA1::P_gpdA_::RFP-H2A::pyr4* |
| Δ*cnaA*Δ*cchA*^rfp-ph-osbp^ | Δ*ku80; pyrG1;* Δ*cnaA::ptrA;* Δ*cchA::hph; AMA1::P_gpdA_::RFP-PH^OSBP^::pyr4* |
| Δ*cnaA*Δ*cchA^vac^* | Δ*ku80; pyrG1;* Δ*cnaA::ptrA;* Δ*cchA::hph; AMA1::P_gpdA_::cpyA-Aeq::pyr4* |
| WT^luc^ | Δ*ku80; pyrG1; P_gpdA_::Luc::pyr4* |
| Δ*cnaA^luc^* | Δ*ku80; pyrG1;* Δ*cnaA::ptrA; P_gpdA_::Luc::pyr4* |
| Δ*cnaA*Δ*cchA^luc^* | Δ*ku80; pyrG1;* Δ*cchA::hph;* Δ*cnaA::ptrA; P_gpdA_::Luc::pyr4* |
| Δ*cnaA^OE::pmcA^* | ZC03*;* Δ*cnaA::ptrA; P_gpdA_::pmcA::hph* |
| Δ*cnaAOE::pmcA^vac^* | Δ*ku80; pyrG1;* Δ*cnaA::ptrA；AMA1::P_gpdA_::cpyA-Aeq::pyr4; P_gpdA_::pmcA::hph* |
| Δ*cnaA OE::pmcA*^PgpdA-gfp-atg8,^*^cpyA-rfp^* | Δ*ku80; pyrG1;* Δ*cnaA::ptrA；AMA1::P_gpdA_::CpyA-rfp::P_gpdA_::GFP-Atg8::pyr4; P_gpdA_::pmcA::hph* |
