## Supplemental Table S2 for "When and why an essential popular mineral—calcium—is an effective antifungal adjuvant against the human pathogen?"

Table S2. All the primers and primer annotations.

| Name | Sequence | Intention |
| --- | --- | --- |
| T7-cnaA-sgRNA5-F | TAATACGACTCACTATAGGGCGCAAGATGGATCAAGCACGTTTTAGAGCTAGAAATAGC | for the DNA template of cnaA-sgRNA3(RNA); for deleting *cnaA* |
| cnaA-ptrA-F | GACGTAATATTCCTTAGTTCACACTGCGCAAGCTAGGAGATCGTCCGCCGATG | for the repair template of deleting *cnaA* |
| cnaA-ptrA-R | CAGATCTTCAGGGCCCAGAGATATAAAACGCAGCCTAGATGGCCTCTTGCATC | for the repair template of deleting *cnaA* |
| cnaA-seq-F | AACCATTATCCACCGGAGGATTG | diagnostic primer for Δ*cnaA* |
| cnaA-seq-R | TGATTCCTTCGGCGCCAAGCATC | diagnostic primer for Δ*cnaA* |
| hph-F | GAATTCCCTTGTATCTCTAC | for *hph* |
| hph-R | TCGAGTGGAGATGTGGAGTG | for *hph* |
| cnaA-not1-F | AAGGGCAATTCGCGGCCCTGATTCTGATAGTAGCACAGCG | for the revertant of *cnaA* mutant |
| cnaA-not1-R | CGAATTGAATTTAGCGGCCCGCATGACAGGTTGAATGCAG | for the revertant of *cnaA* mutant |
| T7-cnaB-sgRNA-F | TAATACGACTCACTATAGGGTTGTTTCAAGGAGAGGACGTTTTAGAGCTAGAAATAGCA | for the DNA template of cnaB-sgRNA |
| cnaB-ptrA-F | ACTCCTCAACCACACACGACCATTTACCAACGCCTAGATGGCCTCTTGCATC | for the repair template of deleting *cnaB* ORF |
| cnaB-ptrA-R | TCAATCACAATCGGTCAGTCCTCTCCTGTCCTAGGAGATCGTCCGCCGATG | for the repair template of deleting *cnaB* ORF |
| cnaB-seq-F | CTCGCCTCTACTGCACGTTC | diagnostic primer for Δ*cnaB* |
| cnaB-seq-R | AACGCCAGGTAGCAAGACTC | diagnostic primer for Δ*cnaB* |
| cnaB-notI-F | AAGGGCCAATTCGCGGCCGTTTCCACCTGTTGTAG | for the revertant of *cnaB* mutant |
| cnaB-notI-R | CGAATTGAATTTAGCGGCCCCTTGGATAGAAGAGATCTGCC | for the revertant of *cnaB* mutant |
| Ama1-BamHI-gpd-F | CGGTTATGCCGTATGGATCCGCATGCGGAGAGACGGACG | for *gpdA* promoter |
| Ama1-BamHI-trpC-R | AATCAGCCTAGCTAGGATCCCATGCATTGCAGATGAGCTG | for *trpC* terminator |
| ClaI-cpyA-F： | CTTTAATCAAGCTTATCGATATGAGAGTTCTTCCAGCTACA | for *cpy*A ORF |
| ClaI-cpyA-R： | TCGAGGTCGACGGTATCGATTTAGAACCATTCACCACCAAGC | for *cpyA* ORF |
| CpyA-linker-R | GGCACCGGCTCCAGCGCCTGCACCAGCTCCGAACCATTCACCACCAAGCC | for *cpyA* |
| Linker-GFP/RFP-F | GGAGCTGGTGCAGGCGCTG | for GFP/RFP |
| Ama1-BamHI-GFP-R | TCAGCCTAGCTAGGATCCTTATTTGTATAGTTCATCCATGC | for GFP |
| Ama1-BamHI-RFP-R | AATCAGCCTAGCTAGGATCCTTAGGCGCCGGTGGAGTGGC | for RFP |
| Linker-Aeq-F | CAGGCGCTGGAGCCGGTGCCATGACCTCCAAGCAGTACT | for Aeq |
| ClaI-pmcA-F | CTTTAATCAAGCTTATCGATATGTCATCGAATCCAAACCAA | for *pmcA* ORF |
| ClaI-pmcA-R | TCGAGGTCGACGGTATCGATCTAGCTTTGTCGCGACTGGC | for *pmcA* ORF |
| RT-pmcA-F | CGATCTCGCTCTAACTCTGC | RT primer for *pmcA* |
| RT-pmcA-R | TAATCTCCGACGAGACGCTC | RT primer for *pmcA* |
| ClaI-pmcB-F | CTTTAATCAAGCTTATCGATATGCTCAATCCCAAGTCGCTC | for pmcB ORF |
| ClaI-pmcB-R | CGAGGTCGACGGTATCGATTTAAGAGCTCGTAATTGTAATGGGA | for *pmcB* ORF |
| RT-pmcB-F | ATGTGACTCGCCTGGACGCTG | RT primer for *pmcB* |
| RT-pmcB-R | CAGGCCACATCATAGACGAG | RT primer for *pmcB* |
| Erg11A-R | CTTGGATGTGTTTTTCGACCGCTTC | for P_gpdA_-Erg11A |
| Erg11A-GFP-F | CGGTCGAAAAACACATCCAAGGGAGCTGGTGCAGGCGCTGG | for GFP |
| Ama1-BamHI-MrsA-F | AATCAGCCTAGCTAGGATCCTTCGTAAACCTACAGCTCCTG | for MrsA |
| MrsA-R | CTCCTGGCGTTTGAAGTAG | for MrsA |
| MrsA-RFP | CTACTTCAAACGCCAGGAGGGAGCTGGTGCAGGCGCTG | for RFP |
| Gpd-R | ATCGATAAGCTTGATTAAAGGTT | for *gpdA* promoter |
| GpdA-RFP-F | GAACCTTTAATCAAGCTTATCGATATGGCCTCCTCCGAGGACGTCA | For RFP-PH^OSBP^ |
| Ama1-BamHI-phosbp-R | AATCAGCCTAGCTAGGATCC TCACGAATTCTTCTTCACAGC | For RFP-PH^OSBP^ |
| Gpd-GFP-F | TTTAATCAAGCTTATCGATATGAGTAAAGGAGAAGAACTTTTCAC | for GFP-Atg8 |
| Ama1-BamHI-Atg8-R | AATCAGCCTAGCTAGGATCCTCAGCAGTCACCGAAAGTGTTC | for GFP-Atg8 |
| Ama1-Bam1-atg8(p)-F | CGGTTATGCCGTATGGATCCCAGCCCACCGTGTAGTAGAG | For *atg8* promoter |
| Atg8-promoter-R | CTTGATAGATAAGGGCGGATAACG | For *atg8* promoter |
| Atg8(p)-GFP-F | GTTATCCGCCCTTATCTATCAAGATGAGTAAAGGAGAAGAACTTTTCAC | for GFP-Atg8 |
| TAA-RFP-R | TTAGGCGCCGGTGGAGTGGC | primer for P_gpdA_-CpyA-RFP |
| RFP-gpdA-F | GCCACTCCACCGGCGCCTAA  GCATGCGGAGAGACGGACG | Primer for P_gpdA_-GFP-Atg8 |
| T7-atg2-F | TAATACGACTCACTATAGGGCAAGAAGTAAGCCATCTGTTTTAGAGCTAGAAATAGCA | for the DNA template of atg2-sgRNA1, for deleting *atg2* |
| Atg2-hph-F: | GAGCCGCTGGTTGGCTCGTTTCGCCCGAGGAATTCCCTTGTATCTCTACACAC | for the repair template of deleting *atg2* |
| Atg2-hph-R: | GAAGGCTCGCTTGACTTTGGCTGCACGGGTGTAGTCTCGAGTGGAGATGTGGAGTGGGC | or the repair template of deleting *atg2* |
| Atg2-seq-F | TGATCATCGCTCGCTATCTGC | diagnostic primer for Δ*atg2* |
| Atg2-seq-R | CTGTGTCTCCAAGATTGTCG | diagnostic primer for Δ*atg2* |
| T7-cchA-sgRNA1-F | TAATACGACTCACTATAGGGTCTGCTTTGCAGCGGGCGTTTTAGAGCTAGAAATAGCA | for the DNA template of cchA-sgRNA1; for deleting cchA |
| cchA-hph-F | TACATTCGGCTAGCGCATTATACCTGCCTACGGAATTCCCTTGTATCTCTACACAC | for the repair template of deleting *cchA* |
| cchA-hph-R | TGCGACATCGACTTCATCATCGAACCCGCCTCGAGTGGAGATGTGGAGTGGGC | for the repair template of deleting *cchA* |
| cchA-seq-F | ACAGCTGCAGTCTGCAATGTC | diagnostic primer for Δ*cchA* |
| cchA-seq-R | GATTGTTGTACATCCTCCACGC | diagnostic primer for Δ*cchA* |
| NOTI-gfp-F | CGAATTGAATTTAGCGGCCGGAGCTGGTGCAGGCGCTGG | for GFP |
| NOTI-gfp-R | AAGGGCAATTCGCGGCCCTATTATTTGTATAGTTCATCCATGCC | for GFP |
| NotI-PgpdA-F | AAGGGCCAATTCGCGGCCGCGCATGCGGAGAGACGGACG | for *gpdA* promoter |
| NotI-TtrpC-R | AATTGAATTTAAGCGGCCGCCATGCATTGCAGATGAGCTG | for *trpC* terminator |
| ptrA-F | CATGGCAGACACTGAAGCAAC | for *ptrA* |
| ptrA-R | TAGCCTAGATGGCCTCTTGCATC | for *ptrA* |
| zero-F | ACTAGTCCTGCAGGTTTAAACGAA | linearizating p-zero-ptrA |
| zero-R | CCCTTTAGTGAGGGTTAATTCTG | linearizating p-zero-ptrA |
| zero-PgpdA-F | TTTAAACCTGCAGGACTAGTGCATGCGGAGAGACGGACG | for *gpdA* promoter |
| PgpdA-R | ATCGATAAGCTTGATTAAAGGTTCT | for *gpdA* promoter |
| PgpdA-GFP-F | AGAACCTTTAATCAAGCTTATCGATATGAGTAAAGGAGAAGAACTTTTCAC | for GFP |
| zero-GFP-R | AATTAACCCTCACTAAAGGGCTGGATCTCGGAGATTTTGTATAG | for GFP |
| CchA-GFP-R | TTGTGGCTATTCGAAGCCATCTGGATCTCGGAGATTTTGTATAG | for P_gpdA_-GFP |
| CchA-F | ATGGCTTCGAATAGCCACAAC | for CchA^N300^ |
| Ama1-BamHI-CchA^N300^-R | AATCAGCCTAGCTAGGATCCTTATTGATCATCAGACATTCCGACAGAC | for CchA^N300^ |
| ClaI-Luc-F | CTTTAATCAAGCTTATCGATATGGAAGATGCCAAAAACATTAAG | for *luc* |
| ClaI-Luc-R | TCGAGGTCGACGGTATCGATTTCTTGGCCTTAATGAGAATCTCG | for *luc* |
| RT-pmcC-F | CTCTGGCCTTAGCAACCGAC | RT primer for *pmcC* |
| RT-pmcC-R | TGAAGATGACGGTGTCGAGC | RT primer for *pmcC* |
| RT-yvcA-F | GCGATCTTCGCTCTGAAGAC | RT primer for *yvcA* |
| RT-yvcA-R | ATCCTCTTCTCCCTGACGAC | RT primer for *yvcA* |
| RT-vcxA-F | GATGACTCTCAACTTCCACATC | RT primer for *vcxA* |
| RT-vcxA-R | GCATCGCCAGTAGGATCATC | RT primer for *vcxA* |
| RT-vcxB-F | GTCATGTTACTGGTGTCAACTGC | RT primer for *vcxB* |
| RT-vcxB-R | TGTCTTTGTCCATGCACCAGC | RT primer for *vcxB* |
| RT-vcxC-F | CACCAAGCTCACTCTGACC | RT primer for *vcxC* |
| RT-vcxC-R | GATTTGATTGTCCCAGGCTTCGTC | RT primer for *vcxC* |
| RT-vcxD-F | GAACCCATCAACGTGTCCAAC | RT primer for *vcxD* |
| RT-vcxD-R | GTAATTCAGTGCAATTCCAACTGG | RT primer for *vcxD* |
| RT-vcxE-F | GTTGGTTCCATTGACGCTCTTAC | RT primer for *vcxE* |
| RT-vcxE-R | CCATCGGCAATGAGATAGTTCAC | RT primer for *vcxE* |
